# Cooperative assembly confers regulatory specificity and long-term genetic circuit stability

**DOI:** 10.1101/2022.05.22.492993

**Authors:** Meghan D. J. Bragdon, Nikit Patel, James Chuang, Ethan Levien, Caleb J. Bashor, Ahmad S. Khalil

## Abstract

In eukaryotes, links in gene regulatory networks are often maintained through cooperative self-assembly between transcriptional regulators (TRs) and DNA cis-regulatory motifs, a strategy widely thought to enable highly specific regulatory connections to be formed between otherwise weakly-interacting, low-specificity molecular components. Here, we directly test whether this regulatory strategy can be used to engineer regulatory specificity in synthetic gene circuits constructed in yeast. We show that circuits composed of artificial zinc-finger TRs can be effectively insulated from aberrant misregulation of the host cell genome by using cooperative multivalent TR assemblies to program circuit connections. As we demonstrate in experiments and mathematical models, assembly-mediated regulatory connections enable mitigation of circuit-driven fitness defects, resulting in genetic and functional stability of circuits in long-term continuous culture. Our naturally-inspired approach offers a simple, generalizable means for building evolutionarily robust gene circuits that can be scaled to a wide range of host organisms and applications.

## INTRODUCTION

In cells, gene regulatory networks integrate and process external and internal information into appropriate gene expression output responses (Levine and Davidson, 2005). Connections in these networks are mediated by the binding of transcriptional regulators (TRs) to DNA cis-regulatory motifs (CRMs) located in upstream proximity to sites of transcriptional initiation. Proper cellular function critically depends on the genome-wide fidelity of these interactions: TRs must functionally recognize gene-associated CRMs with high specificity while avoiding off-target interactions that can result in aberrant misregulation (**Figure 1**). Indeed, there is evidence that native regulatory network fidelity is optimized during evolution, likely through a combination of positive and negative selection processes that, respectively, maximize on-target regulation while minimizing off-target misregulation (Berg et al., 2004; Froula and Francino, 2007; Gera et al., 2022; Hahn et al., 2003; McKeown et al., 2014; Moses et al., 2003; Sengupta et al., 2002; Tsong et al., 2006; Zarrinpar et al., 2003). Disruption of network fidelity due to altered TR specificity or expression levels can lead to a loss in cellular fitness or, in the case of multicellular organisms, to abnormal development or oncogenesis (Gillinder et al., 2017; Lee and Young, 2013).

**Figure 1.**
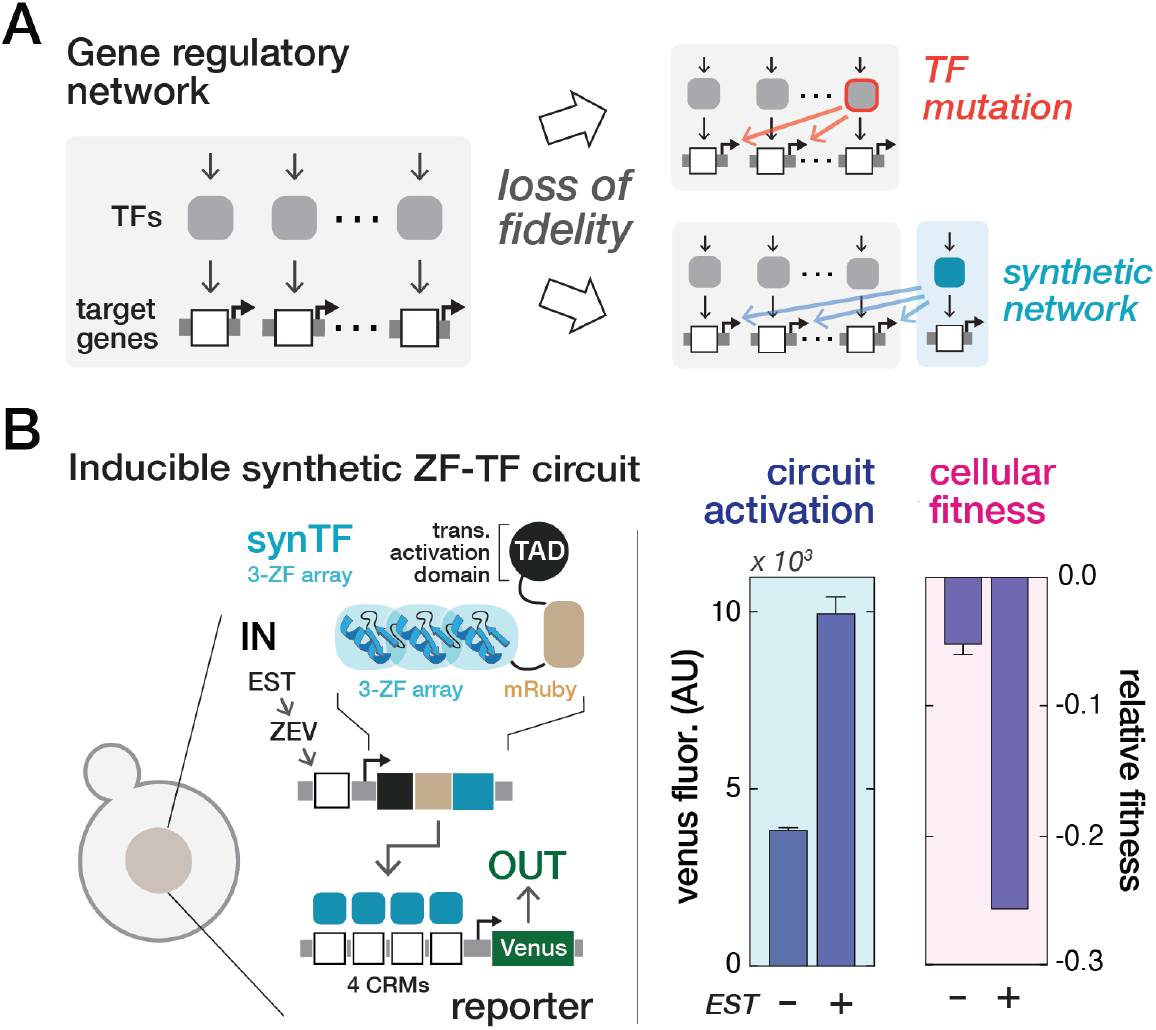
Gene regulatory networks rely on specific molecular interactions between transcriptional regulators and target genes for proper cellular function. **(A)** Cross-reactivity arising from transcription factor (TF) mutations or the introduction of synthetic circuits can drive loss of genome-wide interaction fidelity and disruption of cellular function and fitness. **(B)** Activation of synthetic gene circuits, constructed from a common class of artificial zinc finger (ZF)-based synthetic transcription factors (synTFs), results in observable fitness defects in yeast. The inducible circuit was chromosomally integrated into yeast, induced by addition of ß-estradiol (EST), and circuit activation and cellular fitness were quantified by single-cell flow cytometry for Venus reporter fluorescence and pairwise growth competition, respectively, 36 hours following induction. Bars represent mean values for three biological replicates +/− SD.

Extensive and ongoing investigation into the molecular basis of transcriptional regulation has revealed that strategies employed by cells to maintain network fidelity can vary dramatically across phylogeny (Morgunova and Taipale, 2017; Wunderlich and Mirny, 2009). For example, network connections in prokaryotes are maintained by families of TRs (e.g., helix-turn-helix and winged-helix members) that recognize CRMs via large-footprint, high affinity interactions capable of specifying unique addresses within small sized genomes (10^6^-10^7^ bp) (Browning and Busby, 2004; Madan Babu and Teichmann, 2003; Wunderlich and Mirny, 2009). By contrast, despite possessing much larger genomes (10^7^-10^9^ bp), eukaryotic cells primarily regulate transcription using TRs (e.g., zinc finger and homeobox family TRs) that recognize and weakly bind to short, highly degenerate CRMs that occur at locations scattered throughout the genome (Crocker et al., 2015; Jaeger et al., 2010; Jolma et al., 2013; Lambert et al., 2018; Li et al., 2008; Tanay, 2006; Zhu et al., 2009). How then can network fidelity be established with such low specificity TRs that are incapable of cognate CRM recognition within a complex genome? One explanation is that regulatory connections in eukaryotic networks are established via the cooperative assembly of TRs at closely spaced clusters of CRMs located within enhancer regions (Bell et al., 1988; Friedlander et al., 2016; Morgunova and Taipale, 2017; Panne, 2008; Ptashne, 1988; Todeschini et al., 2014). Under this scheme, CRM proximity strengthens TR binding through weak multivalent interactions between TRs and with associated transcriptional cofactors (Farley et al., 2015; Levine, 2010; Morgunova and Taipale, 2017; Spitz and Furlong, 2012). Thus, because functional regulatory connections are dependent on the cooperative assembly of multiple TRs, eukaryotes can maintain network fidelity despite the potential for genome-wide CRM occurrence (Gao et al., 2018).

Over the last two decades, engineering artificial transcriptional regulatory networks to reprogram cellular behavior has become a major focus for the field of synthetic biology (Cameron et al., 2014; Meng and Ellis, 2020) and has emerged as a powerful approach for the development of cell-based industrial and biomedical technologies (Fischbach et al., 2013; Kitada et al., 2018; Voigt, 2020; Xie and Fussenegger, 2018). These engineered networks, often termed gene circuits, are constructed using TR-CRM interactions that specify links between genes, or couple gene expression outputs to molecular inputs such as small molecules, proteins, or RNA (Bashor and Collins, 2018; Brophy and Voigt, 2014). To date, the predominant focus in the field has been on identifying molecular parts (e.g., engineered TRs and promoters) and validating design strategies that enable the construction of gene circuits with quantitatively precise steady-state and dynamic behavior. Circuits engineered in both prokaryotic and eukaryotic host cells are currently under development for a wide range of applications, including metabolic engineering and cellular therapy. An emerging and critical feature of designing circuits for these real-world applications is their genetic stability (Son et al., 2021). Introducing gene circuits into host cells can impose a fitness cost by creating a metabolic or resource burden, or from expression of a toxic protein product (Borkowski et al., 2016; Ceroni et al., 2015; Gorochowski et al., 2017). Cells harboring mutations that abrogate circuit function can therefore acquire selective growth advantages over those with functionally intact circuits, leading to the progressive loss of circuit-bearing cells from a population. While design strategies for addressing these issues have been described (Ceroni et al., 2018; Gorochowski et al., 2017; Muller et al., 2019; Ng et al., 2019; Riglar et al., 2017), most rely on challenging ad hoc debugging, and generalizable rules for engineering circuit stability remain mostly undefined.

Disruption of transcriptional network fidelity represents another potential source of instability for gene circuits. To date, most circuit engineering efforts have focused on designing TR-CRM interactions to support robust regulatory connections. However, it is seldom investigated whether circuit expression leads to diminished fitness through spurious interactions between circuit TRs and non-cognate CRMs within the host cell genome. Indeed, loss of network fidelity resulting from circuit-associated TR expression may pose an acute challenge for gene circuits engineered in eukaryotic cells, which are often constructed using low-information TRs with potential for off-target misregulation. We recently developed a gene circuit engineering platform in yeast that uses synthetic zinc finger (ZF)-derived transcriptional activators to mediate circuit connectivity (Khalil et al., 2012). As our previously published work demonstrates, this framework can be readily utilized to construct diverse synthetic network connectivity, enabling precise control over circuit dose response as well as programmable Boolean logic and complex temporal behavior (Bashor et al., 2019). In this study, we investigate the genetic stability of circuits engineered using this framework. We show that an observable fitness cost associated with circuit activity is caused by off-target misregulation of host cell transcription, leading to the gradual loss of circuit function across a cell population. In order to restore network fidelity, we draw upon the organization of natural networks as inspiration and test whether cooperative TR assembly can be used as an engineering strategy to create insulated regulatory connections that limit off-target TR binding. As our results show, circuit connections that are functionally dependent on multivalent assemblies can be used to effectively mitigate misregulation and restore fitness, resulting in the long-term stabilization of circuit function.

## RESULTS

Our recently reported synthetic gene circuit engineering platform recapitulates many of the essential design features of native transcriptional regulation in yeast and other eukaryotes. The platform comprises a set of synthetic transcription factors (synTFs) constructed from Cys2-His2 ZFs, the most prevalent and conserved DNA-binding domain across eukaryotes (Lambert et al., 2018; Wolfe et al., 2000). The creation of tunable network linkages using synTFs is facilitated by their modular design; they are composed of 3-finger ZF domain arrays engineered to bind artificial ∼9 bp CRMs that are arranged in clusters upstream of a core promoter. Appending transcriptional activation domains or protein-protein interaction domains to either terminus of the ZF array enables synTFs to, respectively, activate transcription at the core promoter and interact with synTFs bound to adjacent CRMs (**Figure S1A**). The strength of synTF-mediated circuit linkages can be tuned by adjusting molecular parameters such as the number of CRMs and the strength of ZF binding. Furthermore, we created a collection of 20 distinct ZF species with orthogonal CRM specificities that facilitate robust construction of circuit designs featuring multiple synTFs. Our work and that of others has demonstrated the utility of programming synTF circuits for a variety of circuit functions in host cells that span eukaryotic phylogeny, including in therapeutically relevant human cells (Bashor et al., 2019; Donahue et al., 2020; Keung et al., 2014; Khalil et al., 2012; Lohmueller et al., 2012; Newby et al., 2017; Nguyen et al., 202; Park et al., 2019; Zhu et al., 2022a; Zhu et al., 2022b).

Since the CRMs that our synTFs interact with are of a similarly low information content as those of native eukaryotic ZF-TRs (Stewart et al., 2012; Wunderlich and Mirny, 2009), there is a possibility for off-target interactions between synTFs and genomic CRM sites, potentially leading to perturbation of host cell transcriptional network fidelity. While the diminished circuit performance or host fitness that accompany a loss of fidelity may go unobserved during short timescale experiments that test synTF circuit dynamics, it is possible that such phenotypic defects may manifest during longer timescale experiments that involve cell growth over many generations. This possibility motivated us to test whether there are measurable fitness costs associated with expression of synTF circuitry in yeast. We constructed a prototype inducible circuit consisting of a single transcriptional network linkage in which expression of a synTF containing a ZF from our collection (42-10) is under the control of an estradiol (EST)-inducible system to activate expression of a Venus reporter gene (**Figure 1B, S1A, B, Methods**). Following induction with EST, we observed expected reporter activation. However, we also observed a concomitant loss of cellular fitness as measured over 36 h in pairwise growth competition with a reference control strain (**Figure 1B, S1C**). Control experiments confirmed that synTF expression was the source of both circuit activation and the fitness penalty; expressing the combination of ZF with TAD, but not either domain independently, led to a fitness decrease (**Figure S1D ‘high affinity’**). To investigate whether this result was specific to ZF 42-10-derived synTFs, we constructed circuits featuring synTFs containing ZFs from our entire collection (**Figure S1E**). For these 20 synTFs, which contain an average of 97.8 potential genomic binding sites (**Figure S1F**), we observed a consistent pattern of circuit activation and fitness loss, highlighting the generality of this observation.

### Cooperative TR assemblies reduce fitness cost burden while maintaining circuit output

Because of the dependence of the observed growth phenotype on a synTF-TAD fusion, we reasoned that circuit-mediated fitness defects could potentially be the result of altered native gene expression caused by synTFs binding to off-target CRMs throughout the host cell genome. Indeed, analysis of the yeast genome revealed the occurrence of 78 sites that were sequence matches for the core 42-10 CRM, and another 1839 sites containing single base mismatches (**Figure S1F**). Since this abundance of potential off-target sites would make removal via genome editing time consuming and laborious, we considered less complicated engineering strategies that could mitigate fitness cost while maintaining circuit function. The predominant approach for programming network connections in synthetic circuits is based on TRs that have “one-to-one” specificity, a design strategy that mirrors native prokaryotic gene regulation by relying on high-information content, binary TR-CRM recognition to encode regulatory links with genome-wide specificity. On the other hand, regulatory strategies involving cooperative assembly that are common in eukaryotic cells rely on TRs that are individually weakly binding and low information to establish robust, highly specific connections through multivalent association. Since these TRs have molecular characteristics similar to our synTFs, we hypothesized that circuits incorporating regulation by cooperative assembly could potentially be used to engineer synthetic circuits with enhanced fidelity and diminished fitness defects.

To gain insight into molecular strategies for using cooperative synTF assemblies to construct highly specific circuit connections, we constructed a simple thermodynamic-based model of transcription regulation that extends our previous work (Bashor et al., 2019) (**Figures 2A**) (**Methods**). This class of model can offer a simplified first-principles framework for predicting gene expression patterns based on key biophysical properties (e.g. protein-DNA interactions and protein-protein interactions) and can be useful as a guide for understanding synthetic systems in which such properties are design-specified (Bintu et al., 2005; Buchler et al., 2003; Garcia et al., 2011; Gertz et al., 2009). Our model considers the simplified case of a TF that can interact with a CRM at both target synthetic (SYN) and ‘off-target’ native (NAT) loci. As an example, we consider a SYN locus with four tandem CRMs and a NAT locus with a single CRM, where all sites are assumed to be identical (**Figures S2A,B**). CRM binding is governed by TF concentration ([TF]) and its affinity for the CRM (*K*_TF_), and the energy of the cooperative interactions between TRs bound to adjacent sites (*c*) (Buchler et al., 2003). We defined a regulatory specificity score as the difference between transcriptional output at the SYN (*txn*_SYN_) and NAT (*txn*_NAT_) loci (**Figure S2C**), and then plotted this score as a function of [TF], *K*_TF_ and *c* (**Figure 2A right**). This analysis revealed that regulatory specificity improves along an axis defined by lowering affinity for DNA and increasing TF cooperativity, a relationship that remained qualitatively similar for cases containing different numbers of binding sites in both the SYN and NAT loci, as well as different formulations of the regulatory specificity score (**Figure S2D**). This result suggests a design strategy whereby high specificity circuit connections can be obtained by engineering synTFs that enforce complex formation through strong interaction with each other but interact weakly with DNA.

**Figure 2.**
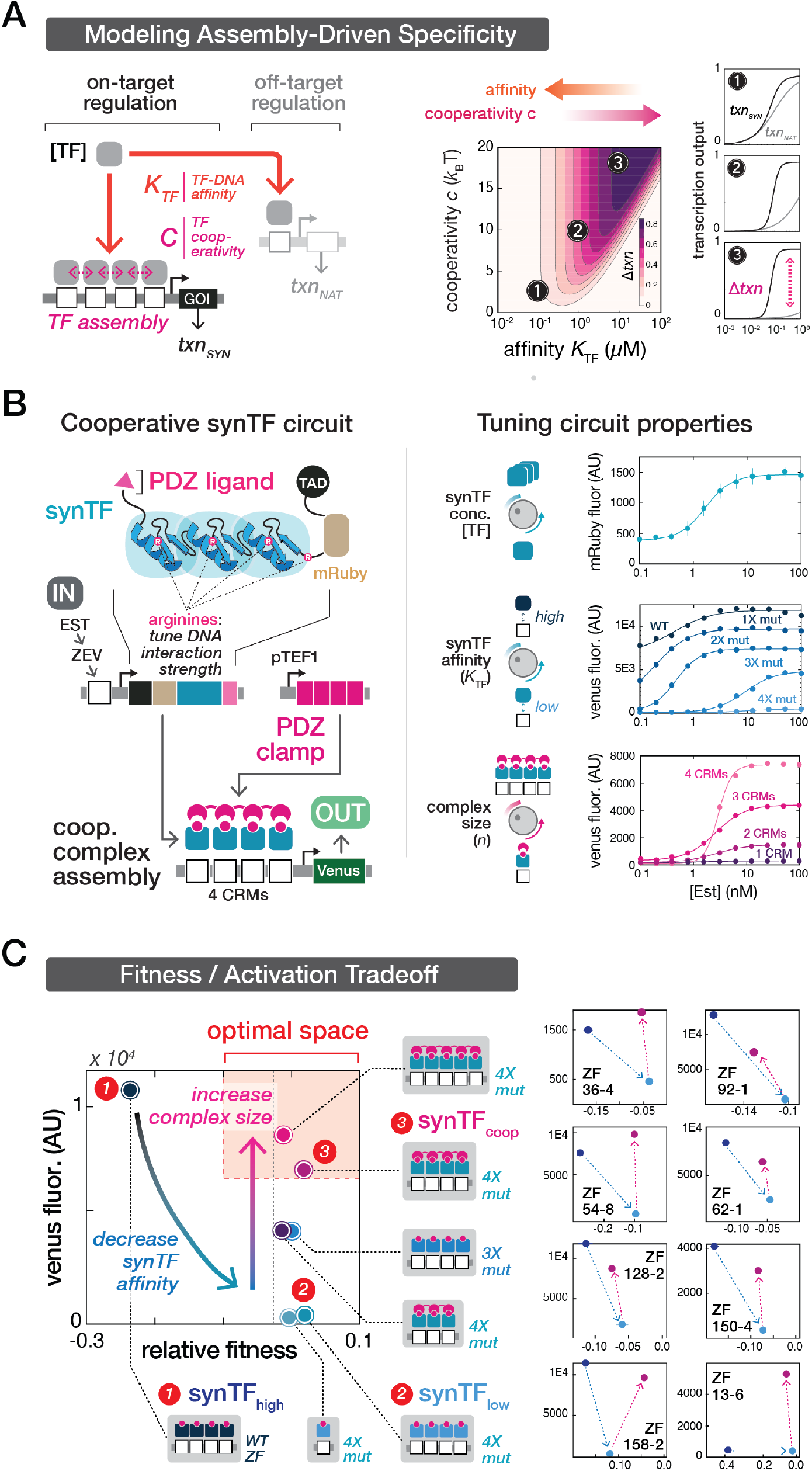
Constructing gene circuits using cooperative assembly minimizes fitness defects while maintaining robust circuit output. (**A**) A simple model reveals a molecular strategy for using cooperative assemblies to engineer regulatory specificity. Left: Thermodynamic-based model of transcription regulation in which a transcriptional regulator (TR) can interact with DNA at both an on-target synthetic circuit (SYN) locus and an off-target native (NAT) locus to regulate gene expression. Binding is governed by the TR affinity for the DNA (*K*_TF_) and the level of cooperativity between TRs bound to adjacent sites (*c*). Right: Regulatory specificity, defined as transcriptional output from the SYN vs. NAT locus, improves as the TR affinity for DNA is lowered and TR cooperativity is increased. (**B**) Experimental platform for constructing circuits composed of cooperative synthetic transcription factor (synTF) assemblies. Left: Inducible circuit architecture. SynTF species, under the control of an estradiol (EST)-inducible system, form multivalent assemblies at a reporter locus to drive gene expression. Complex formation is mediated by a clamp protein. The synTF complex architecture and cooperativity is governed by programmable interaction domains (zinc finger (ZF) and PDZ) and their respective binding partners (CRM and PDZ ligand). Right: Tunable circuit properties. SynTF concentration, DNA affinity, and complex size are rationally tuned by adjusting the EST dose, number of arginine-to-alanine (R→A) mutations at conserved ZF positions that mediate nonspecific DNA interactions, and number of CRM sites, respectively. Points represent mean values for three biological replicates +/− SD. WT, no R→A mutations; 1X mut, 1 R→A mutation; 2X mut, 2 R→A mutations, etc. (**C**) Circuits utilizing cooperative regulatory assemblies of low-affinity synTFs minimize fitness costs while maintaining high circuit output, thus optimizing the fitness-activation tradeoff. Left: Cellular fitness vs. circuit activation for various non-clamp and clamp circuit configurations, all constructed from the same ZF (42-10). Right: Fitness-activation measurements of circuit configurations constructed from different ZF species (with different binding sequences) exhibit similar patterns. Points represent mean values for three biological replicates +/− SD.

We sought to test this strategy experimentally using a framework we recently developed for engineering multivalent synTF assemblies (Bashor et al., 2019). Under this system, interactions between synTFs bound to tandem, core promoter-adjacent CRMs are mediated by a “clamp”: a synthetic protein composed of multiple covalently linked PDZ domains that interact with peptide ligands on synTFs to enable multivalent coordination of their binding to DNA (**Figures 2B left, S3A**) (**Methods**). Modifying the circuit in **Figure 1B** with a constitutively expressed clamp enables us to test relationships between molecular parameters underlying complex assembly and regulatory specificity that were suggested by our model (**Figure 2B**): synTF expression level ([TF]) can be tuned through the addition of EST, while *K*_TF_ for the synTF can be adjusted by introducing alanine mutations (WT - 4x mut) to a set of conserved arginine residues in the ZF array that make non-specific interactions with the DNA phosphate backbone (Elrod-Erickson et al., 1996; Khalil et al., 2012; Pavletich and Pabo, 1991) (**Figure 2B right**). Additionally, *c* can be tuned by varying complex valency (*n*), resulting in altered dose response steepness (**Figures 2B right, S3B**).

We constructed various clamp and non-clamp circuit configurations, tested them for activation and competitive growth rate (**Figure S1D**), and then plotted their mean fluorescence and relative fitness on a two-dimensional “fitness-activation” phenotypic space (**Figure 2C**). The circuit configuration tested in **Figure 1B**—a circuit containing a “high-affinity” (wildtype ZF) synTF (termed the **synTF_high_** circuit) exhibited high reporter activation but low cellular fitness, placing it in the top-left region of the space (**Figure 2C**). Reducing *K*_TR_ by introducing 3 or 4 R→A mutations was sufficient to rescue the fitness, however, this predictably led to significant loss of circuit activation (e.g. see **synTF_low_**: low-affinity 4X mut ZF). Consistent with predictions from our model, we found that circuit activation could be restored via expression of clamp with 4X synTFs in a *n*=4 (synTF_coop_) or 5, with little apparent loss of fitness (**Figure 2C**). Furthermore, we found these effects on fitness and circuit output are not due to differential synTF expression, as low-affinity variants are expressed at equivalent, or even modestly increased, levels compared to synTF_high_ (**Figure S3C**). To determine the generalizability of this result, we tested synTF_high_, synTF_low_, and synTF_coop_ circuit variants for our entire ZF collection and observed the same pattern with all ZFs: a rescue of fitness from synTF_high_ to synTF_low_, and a subsequent improvement of circuit function in synTF_coop_ (**Figures 2C right, S1E**). Our data demonstrate that wiring synTF circuits using cooperative assemblies offers a simple and extensible strategy for optimizing both circuit function and host fitness.

### Cooperative assembly can rescue aberrant gene expression caused by synthetic circuits

To verify that differences in synTF circuit-imposed fitness costs are indeed the result of host cell network misregulation, we performed RNA-sequencing (RNA-seq) to assess host cell transcriptomics following induction of synTF_high_, synTF_low_ and synTF_coop_ circuits (all constructed from ZF 42-10). Biological replicates of each strain demonstrated highly correlated gene expression profiles (**Figure S4A**). We found that synTF_high_ expression led to widespread misregulation of the host transcriptome relative to a control strain with the same genetic background but lacking the synTF (“reporter only”) (**Figure 3A**). Consistent with a general model of TAD-dependent off-target gene activation by synTFs, the majority of misregulated genes were upregulated (182/211) (**Figure 3B**), and more likely to harbor potential synTF binding sites (8/9 bp homology to the CRM) and occur within a 300 bp window upstream of the TSS (14.8% or 27/182 genes) compared to both downregulated (0/29 genes) or unaffected genes (2.8% or 134/4827) (**Figure S4E**).

**Figure 3.**
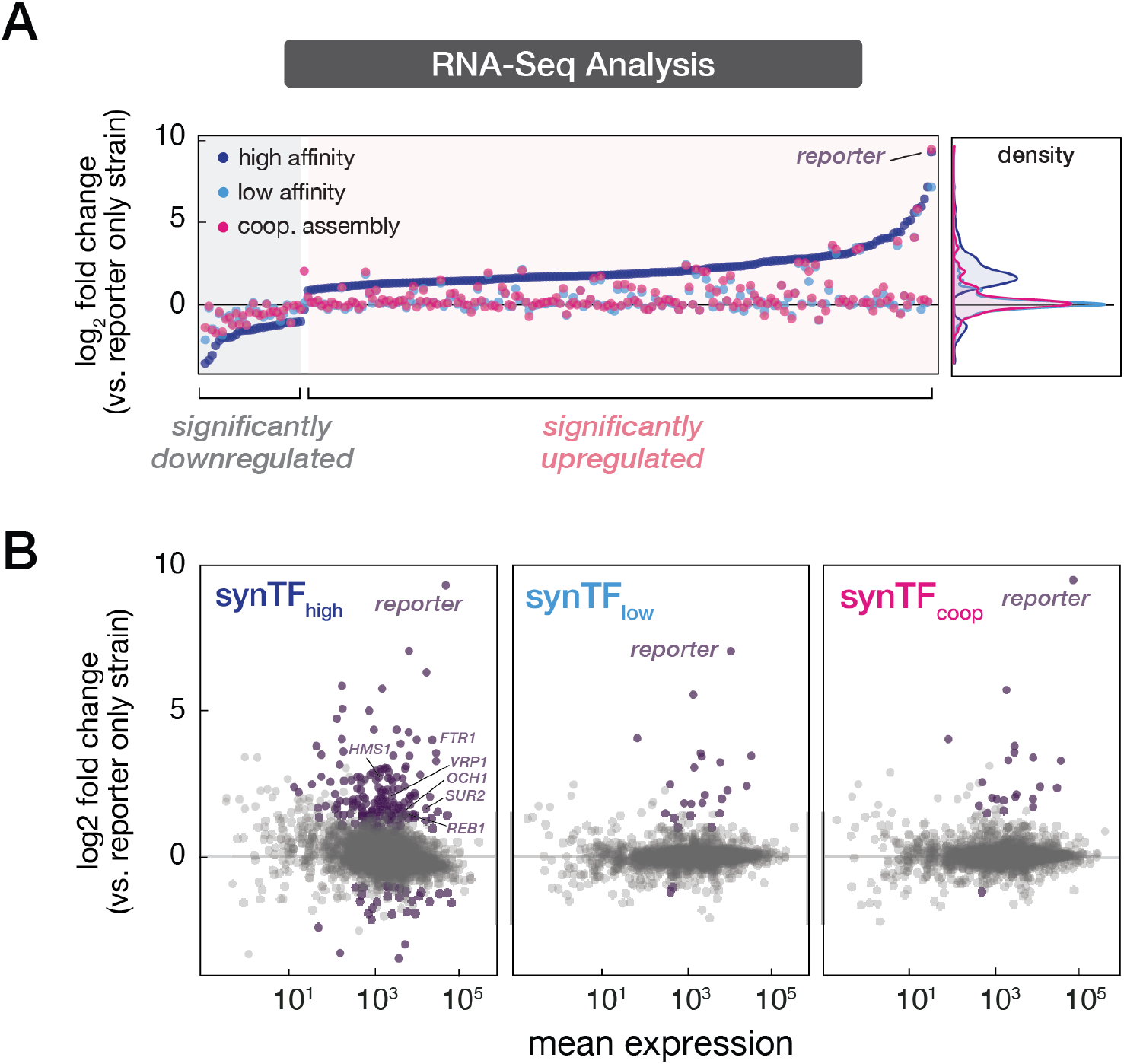
Cooperative assembly rescues aberrant gene expression caused by synthetic circuits. **(A)** Differential gene expression analysis of RNA-seq measurements following the induction of synTF_high_, synTF_low_, and synTF_coop_ circuits. Plotted are genes that are significantly differentially regulated relative to reporter-only control. The synTF_high_ circuit induces a global misregulation of the host transcriptome, including significantly upregulating 182 genes. Gene expression density distributions of synTF_coop_ and synTF_low_ strains are highly similar to one another and cluster tightly around reporter-only background. **(B)** Differential gene expression profiles for synTF_high_, synTF_low_, and synTF_coop_ strains, plotted for all genes. Purple dots denote genes that are significantly differentially regulated vs. reporter-only strain. The reporter was the only differentially regulated gene in synTF_coop_ vs. synTF_low_.

By contrast, transcriptomes of cells harboring the synTF_low_ and synTF_coop_ circuits demonstrated expression profiles that were similar to one another and to the strain background, indicating minimal effect on native transcription (**Figures 3A-B, S4B-D**). As expected, we found that addition of the clamp shows no effect on endogenous gene misregulation compared to the synTF_low_ case (**Figures 3B and S4C**), mirroring the observation that synTF_low_ and synTF_coop_ have similar fitness profiles. In fact, the only gene showing differential regulation between synTF_coop_ and synTF_low_ strains was the fluorescent reporter, with the synTF_coop_ circuit showing comparable expression levels to synTF_high_. Altogether, these results implicate transcriptional network misregulation as the basis of the observed growth defect in the synTF_high_ circuit and demonstrate that this defect can be rescued by tuned-down synTF-CRM interaction affinity in the synTF_low_ and synTF_coop_ circuits.

### Synthetic cooperative assembly reduces off-target binding in the genome

The data revealed by our RNA-seq experiments are consistent with off-target regulation in the host cell genome underlying the fitness cost associated with expression of a high-affinity synTF. To verify that this misregulation is driven by promiscuous synTF binding events, we performed chromatin immunoprecipitation-sequencing (ChIP-seq) analysis of the synTFs across the three circuit strains (synTF_high_, synTF_low_, synTF_coop_), the corresponding strains with synTFs lacking a TAD fusion, and the reporter-only control strain (**Methods**) (**Figure S5A**). Importantly, we spiked in known quantities of *S.pombe*-derived FLAG-tagged DNA, which allowed our data to be normalized to facilitate quantitative comparisons between strains (Doris et al., 2018). In addition to the reporter locus, which showed the expected strong enrichment of synTF binding in both the synTF_high_ and synTF_coop_ strains, for both -/+ TAD conditions (**Figures S5B,C**), we also observed significant enrichment of synTF binding at 23 sites in synTF_high_, 5 in synTF_coop_, and none in synTF_low_ (**Figure 4A**). To evaluate whether these 28 sites could potentially mediate off-target synTF misregulation, we filtered them on the basis of two criteria: (1) whether the site was robust and not a potential pulldown artifact based on its presence for strains both with and without the TAD fusion (Kidder et al., 2011); (2) proximity of the alignment peak (within 700 bp) to a putative synTF CRM, as determined by an independent dataset of quantitated 42-10 binding specificities (Pattanayak et al., 2011) (**Methods**). Satisfyingly, we found that the same 10 sites were solely and independently isolated by both criteria, and were from the synTF_high_ strain (**Figure 4A**). In most cases, these sites contained motifs that were directly under the ChIP peak maxima, with some sites harboring multiple motifs clustered under the peak (**Figure 4B**). Additionally, all of the ten ChIP-seq hits contained top-ranked binding sequences identified from the independent *in vitro* dataset (Pattanayak et al., 2011) (**Figure S5E**). In contrast, none of the 3 robust hits for our synTF_coop_ strain had a correlated motif proximate to a ChIP peak. Furthermore, virtually all of the other motifs that we identified (within 2 kb of peaks) that were not correlated with a likely ChIP-seq binding event were low-ranked ZF binding sequences as determined by the independent *in vitro* experiments. These results provide strong evidence that our ChIP-seq analysis likely identified *bona fide* binding events for synTFs.

**Figure 4.**
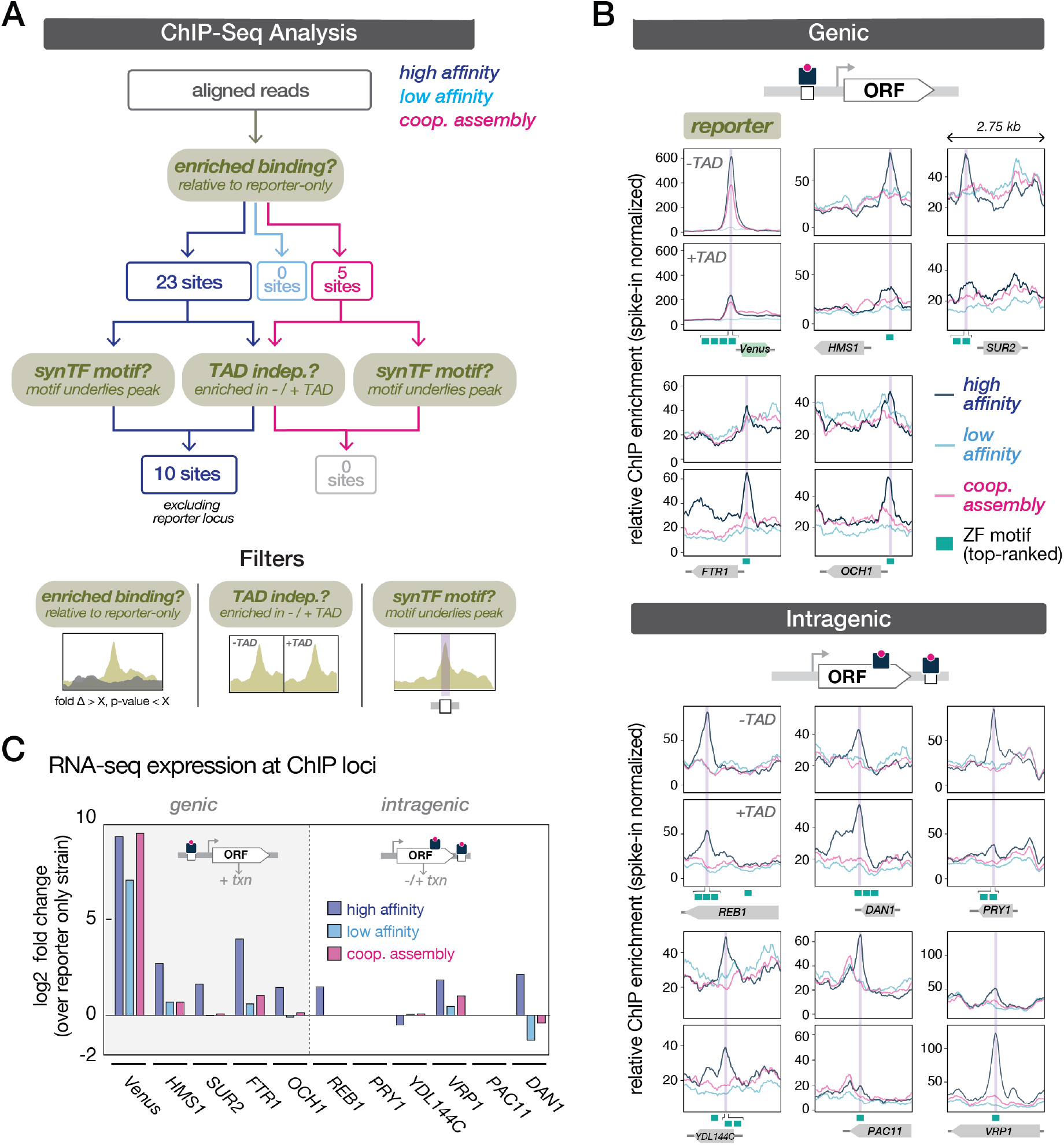
Synthetic cooperative assembly reduces off-target binding in the host genome. (**A**) ChIP-seq analysis pipeline for identifying genome-wide binding events by synTFs. Significant binding enrichment relative to the reporter-only strain was observed at 28 endogenous sites. These hits were subsequently filtered based on two criteria: 1) presence in strains with and without TAD fusion and 2) proximity to synTF motif. This analysis yielded 10 sites enriched in the synTF_high_ strain that were identified as binding events. The synthetic reporter locus was the only enriched site that met these criteria in the synTF_coop_ strain (along with the synTF_high_ strain). (**B**) SynTF ChIP enrichment patterns at the 10 nominated binding sites, classified as genic if located upstream of a gene TSS or intragenic if located within a gene body. Location of top-ranked binding sequences (as determined by *in vitro* studies) are denoted by green boxes and were highly correlated with bound regions. Relative ChIP enrichment was normalized to FLAG-tagged *S. pombe* spike-in DNA that was produced in parallel with the *S. cerevisiae* samples. (**C**) RNA-seq differential expression of genes associated with a synTF ChIP binding event. In general, synTF_high_ genic binding events were associated with higher gene expression, except at the reporter locus where synTF_high_ and synTF_coop_ exhibited similar expression levels. Bars represent the log_2_ transformed fold change in transcription for each strain (synTF_high_, synTF_low_, synTF_coop_) over the reporter-only control at each labeled gene.

To gain further insight into the potential role of the 10 synTF_high_ ChIP-seq-nominated binding events in conferring fitness defects, we plotted alignment peaks from each of the circuit-containing strains atop their corresponding genomic loci (**Figures 4B, S5D**), classifying binding events as genic or intragenic based on the position of the CRM (and associated peak) relative to the nearest gene (Doris et al., 2018). Here, genic denotes a motif located upstream of a gene, where it is more likely to be involved in transcriptional activation, while intragenic denotes a motif located within an ORF, where its effect on gene transcription is *a priori* less clear (*e.g.* positive, negative, no effect). We then plotted the RNA-seq-measured expression changes for the reporter and the ChIP-nominated genes for each of the three circuit strains (relative to reporter-only) and found that transcription of all of the genes associated with genic binding events in the synTF_high_ strain were upregulated relative to the control, whereas those associated with intragenic events showed variable regulation (**Figure 4C**). Importantly, and as expected, the synTF_high_ misregulation patterns were largely rescued in the low-affinity strains, except at the reporter locus which showed comparable activation in synTF_high_ and synTF_coop_ strains. Altogether, these results strongly implicate off-target synTF binding as the likely source of host cell transcriptional misregulation in the strains harboring the synTF_high_ circuit, an effect that is minimized by cooperative synTF assembly in our synTF_coop_ circuit.

### Cooperative synTF regulatory linkages enhance long-term genetic circuit stability

Motivated by the finding that cooperative synTF assemblies can be used to mitigate loss of transcriptional fidelity and the accompanying fitness cost associated with circuit expression, we investigated whether this strategy could also confer long-term circuit stability in continuously growing cultures. To test this, we utilized a customizable, automated bioreactor platform we recently developed, called eVOLVER (Heins et al., 2019; Wong et al., 2018), to perform 5 days-long continuous culture of strains expressing the synTF_high_, synTF_low_, and synTF_coop_ circuit designs along with a reporter-only control (**Figure 5A**). Three biological replicates of each strain (for two different ZFs) were inoculated into separate eVOLVER culture vials, induced with 100 nM β-estradiol, and grown under a turbidostat regime for 130 hours to continuously maintain cultures at a constant density (**Methods**). Growth rates were measured for each culture throughout the experiment, and cultures were periodically sampled to assess circuit output and synTF concentration.

**Figure 5.**
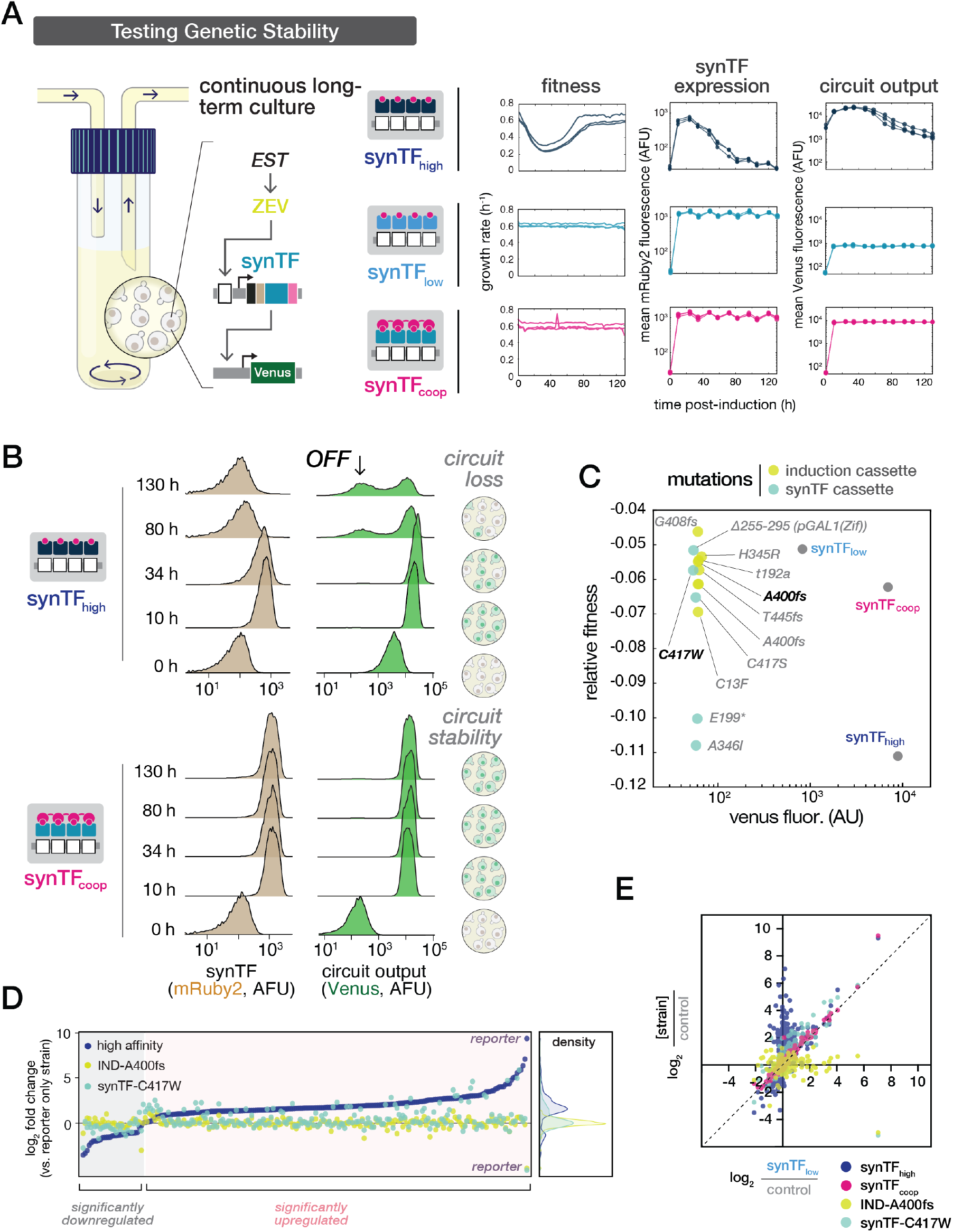
Cooperative regulatory linkages enhance the long-term genetic stability of synthetic circuits. (**A**) Testing long-term stability of synTF circuits in eVOLVER, an automated continuous culture system with real-time measurements of cellular fitness. Individual bioreactor vials were inoculated with three biological replicates of each strain (synTF_high_, synTF_low_, synTF_coop_, reporter-only), induced, and continuously grown in turbidostat culture mode. Samples were periodically taken to measure synTF concentration and circuit output by flow cytometry. Points represent a sample from each of three eVOLVER vials per strain type. (**B**) Single-cell flow cytometry distributions of synTF and circuit reporter expression over the time course of the continuous culture experiment. (**C**) Characterizing circuit genotype mutations selected from the eVOLVER continuous culture experiment. Two classes of mutations were identified from synTF_high_-derived colonies: mutations in the induction cassette (yellow) and synTF cassette (green) (see **Figure S6B**). Each mutation was introduced into a clean synTF_high_ circuit background and quantified for fitness (by growth competition) and reporter expression (by Venus fluorescence). All of the mutations disabled circuit output, while all except two were sufficient to restore fitness to control levels. Points represent mean values for three colonies +/− SD. (**D**) Adaptive circuit-breaking mutations rescue the pattern of gene misregulation induced by the synTF_high_ circuit. RNA-seq differential gene expression analysis for the synTF_high_ circuit and two mutant genotypes: IND-A400fs (induction cassette) and synTF-C417W (synTF cassette). Plotted are genes that are significantly differentially regulated relative to the reporter-only control. (**E**) Correlation of transcriptomes for various circuit genotypes versus the synTF_low_ circuit genotype. Control, reporter-only genotype.

Following circuit induction, we observed a rapid collapse in the growth rate of the synTF_high_ strain, followed by a slow recovery phase over ∼40 hours (**Figure 5A**). This was accompanied by a concomitant decay in the synTF concentration (mRuby) to pre-induction levels, followed by loss of reporter expression (Venus). Single-cell FACS distributions over the time course revealed the gradual emergence of a growing “circuit-off” sub-population, which appeared concurrently with a population-wide loss of synTF-mRuby expression (**Figure 5B top**). In contrast, cultures inoculated with synTF_low_ and synTF_coop_ circuits maintained a growth rate that was similar to the control strain following induction and throughout the duration of the experiment (**Figure 5A**), with synTF concentration and reporter activation remaining unchanged after reaching a post-induction steady state and retaining a sharply defined “circuit-on” population (**Figure 5B bottom**). The same patterns of growth and circuit expression were observed with circuits featuring a different member of our library (13-6) (**Figures S6C,D**).

### Adaptive circuit breaking mutations target synTF expression and function

A plausible explanation for the growth patterns we observed in our eVOLVER experiment is the emergence of adaptive mutations that rescue fitness costs by disabling circuit function, and then ultimately fix within the population by outcompeting cells with intact circuits. To gain insight into whether such mutations could account for our observations, we created a simple computational model designed to simulate populations of cells harboring both functional and broken circuits (**Methods**). The model accounted for the average synTF concentration in each subpopulation of cells while assuming both a fitness cost proportional to the probability of off-target synTF binding, as well as a constant mutation rate capable of disrupting synTF activity and relieving the fitness cost (**Figure S6F**). Consistent with our observed experimental results, our model predicted a decrease in the average culture fitness within 20 hours after induction, followed by a recovery. Furthermore, as the TF-DNA affinity decreases, the model predicts that fitness is improved and the time to recovery increases, while decreasing cooperativity decreases recovery time and reduces fitness (**Figure S6F**). These predictions are consistent with the occurrence of mutations that select against functional synTF expression underlying the observed culture dynamics.

We next experimentally assessed mutational paths to loss of circuit function by analyzing endpoint genotypes from 50 individual colonies from synTF_high_ and synTF_coop_ cultures. No mutations were observed in the circuit genotype from synTF_coop_-derived colonies, while we found a total of 27 mutated synTF_high_-derived colonies, with mutations occurring within the induction cassette (10 distinct) and in the synTF cassette (5 distinct) (**Figure 5C**). Mutations were found in each component of the induction cassette, with one TAD residue (A400fs) targeted in more than a third of all of the colonies (**Figure S6B**). Distinct mutations in the synTF cassette were found in the promoter as well as the coding sequence, with two mutations (C417S and C417W) found in the same cysteine of the synTF Cys2His2 ZF backbone, suggesting that disrupting the ability of the synTF to bind DNA is sufficient for fitness rescue and corresponds with the loss of reporter expression in the mutated strains. To confirm that these circuit mutations drive fitness recovery, we tested the effects of each individually in a clean synTF_high_ background (**Figure 5C**). All of the mutations were shown to disable circuit output, while all but two restored fitness to control levels.

We verified that the mutations were selected for their ability to restore loss of fitness through rescue of host gene network misregulation by performing RNA-seq analysis on two circuit mutants, one from each class: A400fs in the induction cassette (IND-A400fs) and C417W in the synTF cassette (synTF-C417W). We found that either mutation was sufficient to mostly rescue the pattern of gene misregulation induced by synTF_high_ (**Figure 5D**). Interestingly, the transcriptomic profile of the IND-A400fs mutant showed no significant gene misregulation over the reporter-only control, while the profile of the synTF-C417W mutant circuit showed similarity to synTF_low_ and synTF_coop_ strains (**Figure 5E**). These results reinforce functional synTF expression as the basis for synTF_high_ circuit instability and, furthermore, indicate that our engineering strategy for rescuing this fitness defect by lowering synTF affinity recapitulates the growth phenotype of adaptive circuit-breaking mutations.

## DISCUSSION

Our results show that cooperative TR assemblies can be used to engineer highly specific regulatory connections in synthetic gene circuits, offering a means for enhancing circuit performance and minimizing circuit-imposed fitness costs in eukaryotic cells. We initially observed that expression of synthetic gene circuits constructed from synTFs—a widely used class of ZF-based TRs—results in observable growth defects due to misregulation of the native transcriptional network in yeast. Using long-term continuous culture experiments, we demonstrated that these fitness costs drive the gradual loss of circuits from the population as adaptive mutants with abrogated circuit function acquire a selective growth advantage over circuit bearing cells. In agreement with simple models of gene regulation and evolutionary dynamics, we found that network fidelity and host cell fitness could be restored, and circuits stabilized, by engineering cooperative complexes that render circuit connections functionally dependent on multivalent assembly of weakly interacting synTFs. Our work demonstrates that this naturally inspired design strategy can be used to effectively insulate synthetic circuits from the host transcriptional network and could enable the rapid development of circuits with enhanced stability against evolutionary pressures (**Figure 6**).

**Figure 6.**
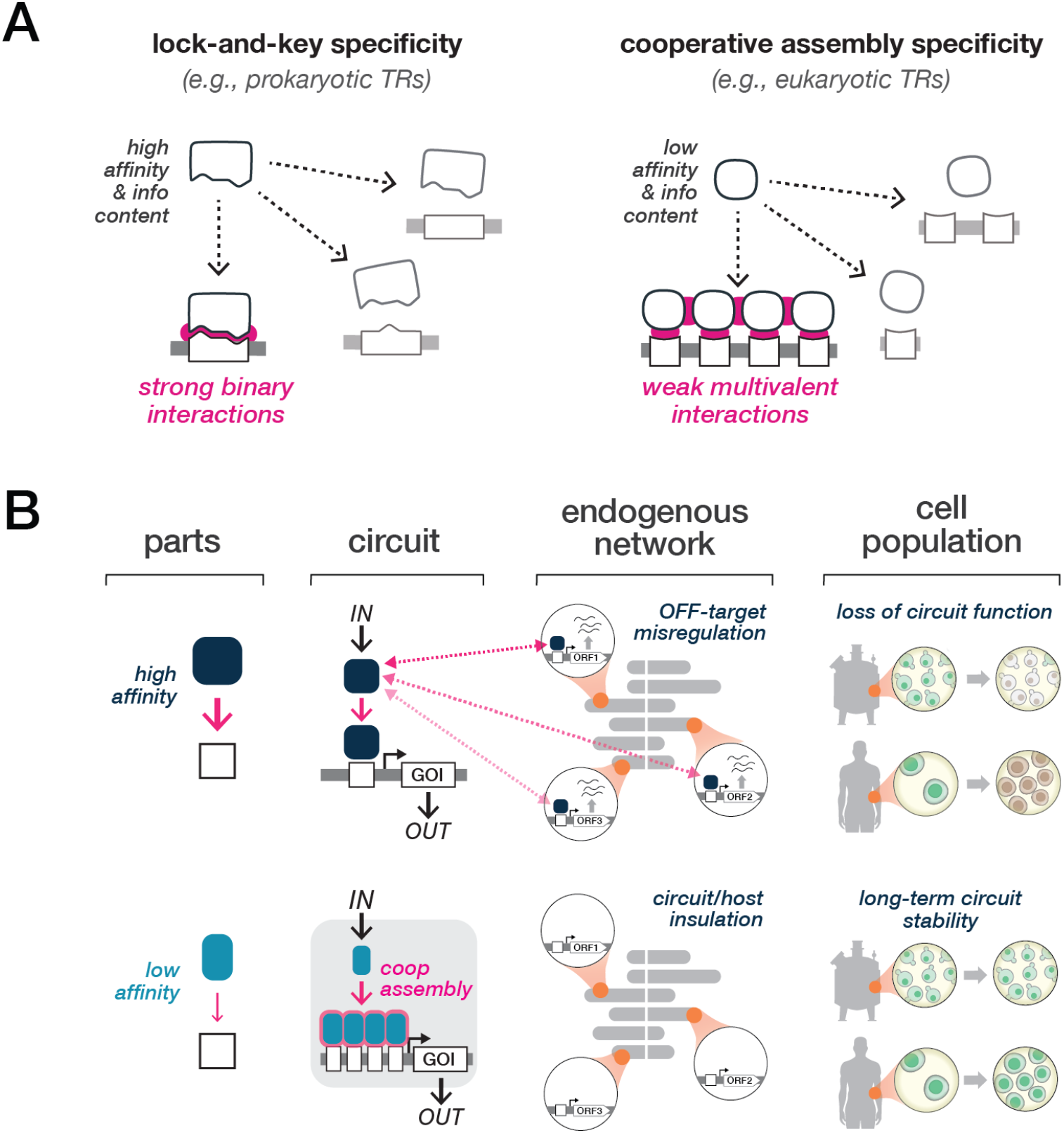
A naturally-inspired strategy to engineer regulatory specificity and long-term genetic circuit stability. (**A**) Strategies employed by cells to maintain regulatory network specificity. One strategy, common to prokaryotic transcriptional networks, uses high affinity, large-footprint interactions between TRs and CRMs in small genomes (lock-and-key specificity). A second strategy, common to gene regulatory networks of higher-order organisms, uses multivalent associations between weakly-interacting, low-specificity components (cooperative assembly specificity). (**B**) Constructing circuit connections using cooperative assembly can effectively insulate synthetic circuits from aberrant misregulation of the host genome to promote long-term stabilization of circuit function.

In recent years, numerous studies have revealed that synthetic circuits are susceptible to unintended interactions with endogenous cellular processes (Borkowski et al., 2016; Ceroni et al., 2015; Gorochowski et al., 2017). These interactions generally impede circuit function, though in some cases they have been shown to serendipitously support it (Cookson et al., 2009; Stricker et al., 2008). Thus, examining the interface between synthetic circuits and the host, and developing strategies to functionally insulate circuits from the host cell have become central objectives in the field of synthetic biology (Nandagopal and Elowitz, 2011; Purnick and Weiss, 2009). Recent studies characterizing synthetic circuits in *E. coli* have established that unintended circuit-host coupling can arise when competition for cellular resources leads to circuit-imposed burden (Borkowski et al., 2016; Gorochowski et al., 2017). These observations have motivated the development of circuit insulation strategies that include: (1) prevention or delay of the emergence of mutants by either integrating circuits, engineering hosts with lower background mutation rate, or maintaining a reduced population size; (2) reducing fitness advantage of circuit mutants by coupling circuit production to an essential gene; (3) using stress response profiling to individually “debug” circuit components, as well as burden reporters that actively modulate circuit activity via feedback control (Ceroni et al., 2015; Ceroni et al., 2018; Gorochowski et al., 2017; Rugbjerg et al., 2018; Sleight et al., 2010).

In this study, we offer evidence that transcriptional misregulation resulting from off-target genomic binding constitutes another class of fitness-reducing circuit-host interaction—one that is potentially a primary source of disruption to circuit function in eukaryotic host cells due to their genomic complexity (**Figure 6B**). Results from our RNA-seq and ChIP-seq experiments provide evidence that the misregulation of host transcription caused by synTF circuits is likely the result of interactions with a select subset of genomic CRMs located primarily, but not exclusively, adjacent to sites of native gene transcriptional initiation. Critically, our data argue that the CRMs responsible for misregulation could not be predicted *a priori*, which motivates the broader question: what are design strategies that could give researchers the ability to rapidly construct new synthetic circuits that confer transcriptional fidelity, given that prediction of off-target TR interactions is complicated by many factors (e.g. regulatory context, chromatin architecture, and cell type) and that relatively few synthetic biology tools are available for optimizing circuits in eukaryotes?

Our solution of using cooperative synTF assemblies to address this challenge draws inspiration from natural eukaryotic transcriptional networks, where combinatorial and cooperative regulation by TFs has been proposed as a key mechanism for the robust and specific rewiring of transcriptional circuitry over evolutionary time (Sorrells et al., 2018; Stefflova et al., 2013; Tuch et al., 2008). This solution is thought to be facilitated by the distribution of binding energy among multiple protein-DNA and protein-protein interactions, which accommodate mutational strengthening or weakening of individual interactions with minimal loss of regulatory robustness (Sorrells et al., 2015). Further, this drift enables the formation and stabilization of new TR-TR and TR-CRM interactions that can facilitate assembly of novel complexes and establish new regulatory connections (Sorrells et al., 2018). Our present work highlights an additional role for cooperative TR assembly as a mechanism to maintain transcriptional network fidelity. Extensive studies have revealed that eukaryotic TRs tend to bind CRMs that are overwhelmingly too short and degenerate to specify unique addresses in the large genome (Tanay, 2006; Wunderlich and Mirny, 2009) (**Figure 6A**). This “specificity paradox” is solved by employing clusters of low affinity binding sites, making specificity and regulatory robustness dependent on the collective action of multiple TRs (Crocker et al., 2015). Indeed, CRMs are often shorter and further from consensus in promoter regions regulated by multiple TRs (Bilu and Barkai, 2005), while frequently interacting TR pairs have been shown to generate composite motifs with unique binding specificity (Jolma et al., 2013; Jolma et al., 2015). Collectively, these observations suggest that by relaxing the importance of any single interaction within a complex, individual TR-DNA interactions are less likely to be functional and deleterious upon the likely appearance of spurious binding sites in a large genome—a strategy that amounts to optimizing the “hub” rather than individually addressing the “spokes”.

From a synthetic biology perspective, our work demonstrates that programming cooperative assembly is a robust, generalizable design strategy for programming insulated synthetic gene circuitry that minimizes cycles of *ad hoc* design (**Figure 6B**). Unlike prevailing strategies that rely on sophisticated biomolecular engineering to develop bespoke, highly specialized components to create regulatory connections, circuits that employ cooperative assemblies can be constructed from existing parts by weakening their interaction affinity and engineering cooperative interactions between combinations of components. This approach requires no *a priori* knowledge of binding and misregulation profiles and, furthermore, minimizes the need to fine-tune regulation of expression levels to manage component toxicity. In addition to simplifying circuit design, engineering cooperative assemblies may provide useful and complementary angles to examine design principles governing how specificity is encoded in natural regulatory systems (Crocker and Ilsley, 2017; Nandagopal and Elowitz, 2011). Cooperative circuits enable systematic studies that are decoupled from crosstalk with native components and regulation of essential processes like differentiation and development.

Finally, because our approach offers a potential means for engineering gene circuit stability, it could prove impactful in biotechnology applications that demand maintenance of circuit function over many generations (**Figure 6B**). For example, in metabolic engineering, strains harboring circuit-controlled biosynthesis pathways must maintain function when they reach bioreactor capacity during growth phases (Lee and Kim, 2015; Wehrs et al., 2019). Similarly, cell-based therapy applications typically require expansion of genetically engineered cells to achieve products that are sufficiently large for patient dosing. In both cases, any burden imposed though circuit-host interactions would not only slow production but could potentially give rise to circuit-deficient subpopulations. In the case of metabolic engineering this might result in uncontrolled or early activation of metabolic pathways that lower yield, while in cell therapy applications, potential effects on product potency and purity could diminish both the safety and efficacy of a treatment. While post-expansion induction of circuits using exogenously activated transcriptional regulatory switches offers one potential solution, the opportunity for misregulation still exits, and the requirement to add an inducer molecule imposes an additional cost on the process. By relieving circuit burden through regulatory insulation, our approach offers a solution to both of these issues that can be applied to existing circuit design strategies by engineering part interactions to accommodate regulatory assemblies. Finally, it is possible that the strategies we developed here could be translated more broadly to genome-manipulating technologies, such as genome and epigenome editing tools, to enhance specificity and enable allele-specific manipulation.

## ACKNOWLEDGEMENT

We thank members of the Khalil laboratory, Jané Kondev, and Zeba Wunderlich for helpful discussions. We are grateful to Fred Winston and his laboratory for providing FLAG-tagged *S. pombe* and technical advice on ChIP-seq experiments. This work was supported by NIH grants R01EB029483 and R01EB027793, NSF grants CCF-2027045 and EF-1921677, ONR grant N00014-21-1-4006, and DoD Vannevar Bush Faculty Fellowship N00014-20-1-2825.

## Author contributions

N.P., C.J.B, and A.S.K. conceived the study. M.D.B., N.P., C.J.B., and A.S.K. designed the study. M.D.B. and N.P. designed and generated all genetic constructs and strains. M.D.B. performed all experiments and analyzed data with N.P. N.P. developed the thermodynamic model of transcription regulation. E.L. developed the evolutionary dynamics model. J.C. assisted with optimization of the RNA-seq and ChIP-seq protocols and performed all the genomics analyses. C.J.B. and A.S.K. supervised the study. M.D.B., C.J.B., and A.S.K. drafted the manuscript with input from all authors.

## METHODS

### Strains

The background strain used for all experiments in this study was *S. cerevisiae* YPH500 (α, *ura3-52, lys2-801, ade2-101, trp1, his3, leu21*) (Stratagene). Strains were constructed by sequential plasmid transformations using standard lithium acetate-based transformation techniques and growth on selective minimal media (Sunrise Science Products), using the *URA3*, *LEU2,* and hygromycin B phosphotransferase (*HPH*, integrated into the *HO* locus) genes as selectable markers. Genotypes for experimentally tested strains are listed in **Table S2**. Experimental replicates comprised distinct colonies picked from a transformation plate following construct integration and selection.

### Cloning and plasmid construction

Plasmid constructs used in this study are listed in **Table S1** and their designs described in **Figures S1 and S3**. All plasmids in this study were constructed using Golden Gate Assembly (Engler et al., 2008) and formatted with the Yeast MoClo Toolkit (Lee et al., 2015) (Addgene Kit #1000000061). ORFs encoding previously described zinc finger and clamp proteins (Bashor et al., 2019; Khalil et al., 2012) were codon optimized for yeast, adapted for Golden Gate assembly, and synthesized (IDT). BsmBI, T7 DNA Ligase, and T4 DNA Ligase Buffer (NEB) were used to construct Level 0 and Level 2 plasmids. The Golden Gate Assembly Master Mix BsaI-HF v1 and v2 (NEB) was used to construct Level 1 plasmids.

### Flow cytometry

Yeast colonies were picked from plates and cultured overnight in 2 mL liquid synthetic defined (SD) media with appropriate auxotrophic dropouts (Sunrise Science Products). Cultures were diluted 1:50 into 500 µL of synthetic complete (SC) media and grown for 7 h at 30°C in the presence or absence of inducer (ß-estradiol). Prior to flow cytometry reading, cells were diluted 1:20 into 200 µL of PBS treated with 20 µg/mL cyclohexamide to inhibit protein synthesis, and stored at 25°C, in the dark, for 1 h to allow for complete fluorophore maturation. Plates were then stored at 4°C overnight. Typically, 10,000 events were acquired using an Attune Nxt Flow Cytometer equipped with a high throughput autosampler (Thermo Fisher Scientific), and data was processed using FlowJo (Treestar Software). Events were gated by forward and side scatter, and geometric means of the fluorescence distributions were calculated. Flow cytometer laser/filter configurations used in this study were mVenus (488 nm, 574/26) and mRuby (561 nm, 620/15).

### Fitness assay

We adapted a previously described fitness assay based on competitive growth between a “reference” strain and “query” strain (Kryazhimskiy et al., 2014). A single colony for a reference strain constitutively expressing (*pTDH3*) an mTurquoise reporter was grown in 2 mL SDC overnight with shaking at 30°C, then diluted 1:100 in 20 mL SDC and grown overnight with shaking at 30°C. Single colonies for three biological replicates for each query strain expressing synTF circuits or the reporter only were each cultured overnight in 500 µL SD -ura/-leu media in 96 well plates. The reference strain culture was diluted 1:50 into 500 µL of SDC in 96 well plates in the presence or absence of inducer (1 µM ß-estradiol) across four 96 well culture plates, and each query strain was added to the reference strain-containing wells at 1:50 and mixed. A 10 µL sample was immediately sampled from each well and fixed in PBS + 20 µg/mL cycloheximide to obtain a *t_0_* measurement of the cocultures prior to induction. The cocultures were then diluted 1:50 into SDC with or without inducer every 12 h and samples were isolated at 16.5 h (*t_1_*) and 36 h (*t_2_*) corresponding to ∼7 and 15 generations, respectively, and fixed in PBS + 20 µg/mL cyclohexamide for flow cytometry analysis. We determined the relative abundances of reference and query strains at *t_0_*, *t_1_*, and *t_2_* for each coculture. Abundance was derived from the fraction of cells in each well expressing the mTurquoise reporter (reference). Fitness was computed for each query strain by calculating changes in abundance from *t_0_* to the experimental endpoint, *t_2_*:

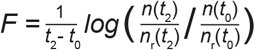

where *n* (*t*) and *n_r_* (*t*) are the cell counts for the query and reference strains, respectively, at time *t* after coculturing.

### Chromatin immunoprecipitation sequencing (ChIP-seq)

#### Preparation, immunoprecipitation, and sequencing

250 mL flasks of SDC were inoculated with overnight cultures and grown for 1 h before induction with 100 nM ß-estradiol, then grown for an additional 8 h to an OD600 of 0.525-0.625. All cultures were diluted to OD600 0.525, then cells were crosslinked with 1% formaldehyde for 9 min at 30°C with shaking. Fixation was quenched with a final concentration of 125 mM glycine (EMD 4840 OmniPur) for 10 min at 30°C with shaking. Cells were pelleted for 10 min at 4°C at 3000 RPM (Haraeus Multifuge X3R), washed twice with ice-cold TE (Tris-HCl, EDTA), transferred to 4 bead-beater tubes/strain and frozen at −80°C. Cell pellets were resuspended in 400 µL ice cold lysis buffer (50 mM HEPES, 140 mM NaCl, 1mM ethylenediaminetetraacetic acid, 1% Triton X-100, 0.1% Na-Deoxycholate, 1 mM phenylmethylsulfonyl fluoride, 200 µL Roche cOmplete protease inhibitors). 0.5 mm diameter glass beads were added to 1 mm below the meniscus. Cells were lysed by bead beating on a MagNA Lyser (Roche) three times for 45 s each at 4500 RPM with 2 min rests at 4°C. Lysate was collected by puncturing the tube with a 21G needle and centrifugation at 2000 g for 2 min into a 2 mL microtube. The pellet was resuspended in lysis buffer, then sonicated for 6 pulses using a probe sonicator (Fisher Scientific FB120) for 20 s at 25% amplitude with 120 s intervening rests on ice, achieving a range of 150-1500 bp DNA fragments. Cell debris was pelleted by centrifugation at max speed for 15 min at 4°C.

FLAG-tagged *S. pombe* (generously provided by the Winston Lab, FWP567) was used as a spike-in control and was prepared similarly to the *S. cerevisiae* cultures with a few modifications: grown in 250 mL YES media to OD600 0.65, split into 5 tubes, underwent 4 lysis steps on the bead beater and 5 sonication steps. The supernatant from the 4 preps of each strain (5 preps of *S. pombe*) were mixed together in a new low retention tube (Thermo Fisher Scientific 02-681-320). To determine DNA concentration, 50 µL samples from each strain were isolated. Samples were brought up to 200 µL with elution buffer, then incubated with 50 µg of RNAse A (Thermo Fisher Scientific) at 37°C for 30 min to remove RNA. Then 100 µg of Proteinase K (Thermo Fisher Scientific) was added and samples were incubated overnight (∼16 h) at 60°C to degrade proteins and reverse crosslinks. Samples were then purified with the ChIP DNA Clean and Concentrator kit (Zymo Research), eluted with 100 µL water and concentrations were determined by Qubit 4 Fluorometer (Thermo Fisher Scientific). 50 µL were brought to 13.5 ng/µL concentration and split into 4 separate tubes, then diluted to 1 mL in lysis buffer. Input samples were concurrently isolated at 10% of the DNA concentration for the IP samples and brought to 100 uL lysis buffer. 1 ug anti-FLAG (Sigma F1804) was added to each IP sample. The prepared (lysed and sonicated) FLAG-tagged *S. pombe* chromatin was added as a spike-in control to 10% of the sample DNA concentration for all IP and input samples. Input samples were stored at 4°C and IP samples rotated overnight at 4°C.

30 µL Dynabeads Protein G (10004D, Thermo Fisher Scientific) per culture was added to a low retention tube and washed 3 times with 1 mL ice cold lysis buffer. Dynabeads were resuspended in 100 µL lysis buffer per culture. 100 µL of Dynabead solution was added to each antibody-pulldown sample and incubated at 4°C for 4 h while rotating. Dynabeads were washed at room temperature on a magnet (twice with 1 mL lysis buffer, twice with 1 mL lysis buffer/500 mM NaCl, twice with 10 mM TrisHCl-pH8/250 mM LiCl/0.5% NP-40/0.5% sodium deoxycholate/1 mM EDTA, and once with 1 mL TE). Bound material was eluted by adding 200 µL of 50 mM Tris-HCl ph8/10 mM EDTA/1% SDS and incubating at 65°C for 30 min. A second elution with the same buffer was combined with the first and tubes were incubated at 65°C overnight to reverse crosslinks. Input samples were brought to 400 µL elution buffer and stored at 65°C with antibody-pulldown samples. 50 µg of RNase A was added to each pulldown and input sample and incubated at 37°C for 30 min. 100 µg Proteinase K was added to each sample, then incubated at 55°C for 4 h. DNA was purified with the ChIP DNA Clean and Concentrator kit (Zymo Research): 4 preps/strain were concentrated into two preps in two columns and eluted with 25 µL of water for IP samples/column (50 µL total) or 200 µL water for input samples and stored at −80°C.

Sample concentrations were measured with the Qubit and 38 µL of each sample was submitted to the Tufts Genomics Core for TruSeq ChIP library preparation (Illumina). Tufts Genomics Core subsequently sequenced all samples, paired end, on a NextSeq 550 (Illumina) to 75 bp.

#### ChIP-seq analysis

ChIP-seq data analyses were performed using the Snakemake workflow management system (Koster and Rahmann, 2012), with code available on GitHub (github.com/khalillab/coop-TF-specificity). Additionally, an archive containing code and raw data suitable for reproducing all ChIP- and RNA-seq analyses is being submitted to Zenodo.

#### ChIP-seq library processing

Adapter removal and 3’ quality trimming of paired-end reads was performed using cutadapt (http://journal.embnet.org/index.php/embnetjournal/article/view/200). Reads were aligned using Bowtie2 (Langmead and Salzberg, 2012) to a combined genome consisting of *S. pombe* genome ASM294v2 concatenated with *S. cerevisiae* genome build R64-2-1 modified to include the mVenus reporter at the URA3 locus. Correctly paired uniquely mapping reads mapping to *S. cerevisiae* were selected using SAMtools (Li et al., 2009). Coverage of fragments and fragment midpoints were generated using SAMtools (Li et al., 2009) and bedtools (Quinlan and Hall, 2010), and normalized to the number of fragments in the library. Quality statistics of raw, cleaned, non-aligning, and correctly paired mapping reads were assessed using FastQC (https://www.bioinformatics.babraham.ac.uk/projects/fastqc/).

#### Transcription factor ChIP-seq peak calling

Transcription factor peak calling was performed for each strain by calling peaks in each replicate using MACS2 (Zhang et al., 2008), followed by filtering for reproducibility among replicates by the Irreproducible Discovery Rate (IDR) method (https://doi.org/10.1214/11-AOAS466). The size of the small and large local regions used by MACS2 to model expected counts were set to 500 and 2000 bp, respectively, and the IDR threshold was set to 0.01.

#### Transcription factor ChIP-seq differential binding analysis

For transcription factor ChIP-seq differential binding analysis, transcription factor peaks were called as described above. A non-redundant list of peaks called in the strains being compared was generated using bedtools (Quinlan and Hall, 2010), and the counts of fragment midpoints from both input and IP samples over these peaks were used as the input to a differential binding analysis with DESeq2 (Love et al., 2014), in which the linear model coefficient extracted represents the change in IP/input enrichment in the query strain versus the control strain. We investigated a set of 132 peaks as candidates for specific binding in any of the synTF_high_, synTF_low_, or synTF_coop_ strains over the reporter-only control strain, at a false discovery rate of 0.05.

### Chromatin immunoprecipitation quantitative PCR (ChIP-qPCR)

Samples were prepared as described for ChIP-seq. qPCR was performed on a LightCycler 480 Instrument II (Roche) with LightCycler 480 SYBR Green I Master Kit (Roche) according to manufacturer’s instructions. A total reaction volume of 10 µL (2 µL of 1:50 dilution of input DNA or 1:20 dilution of IP DNA, 0.5 µM of forward primer, 0.5 µM of reverse primer, 5 µL of 2X SYBR Green Master Mix), using the following cycle conditions: (i) pre-incubation: 95°C for 10 min; (ii) amplification (45 cycles): 95°C for 10 s, 57°C for 20 s, 72°C for 8 s; (iii) melting curve: 95°C for 5 s, 65°C for 1 min, 97°C at ramp rate 0.11C/s; (iv) cooling: 40°C for 10 s. PCR primer sequences were designed to flank the cis-regulatory motifs (CRMs) at the synthetic promoter: gcgatcacagacattaacccacag; tggcggatctgggatccga. Fold enrichment over the reporter-only control strain was then computed from the resulting qPCR Ct values using the ΔΔCt method.

### RNA Sequencing (RNA-seq)

#### Preparation and sequencing

RNA-seq measurements were performed on two biological replicates per strain type. Our results were reproduced with a technical replicate for each biological replicate in two separate experiments, aside from synTF_high_, for which we reported on two biological replicates and a single technical replicate. Total RNA was purified from ∼5×10^7^ cells following the “Purification of Total RNA” from the “Yeast Mechanical Disruption” protocol in the RNAeasy Plus Mini Kit handbook: 50 mL of cells from an overnight culture were induced with *β*-estradiol and cultured for 7 h in a 30°C shaking incubator. Cells were brought to the same concentration, spun down for 5 min at 1000 RCF at 4°C, liquid was removed and the pellets were resuspended in 600 µL RLT buffer + *β*-mercaptoethanol. ∼600 µL of 0.5 mm diameter glass beads were added and cells were lysed by bead beating on a MagNA Lyser (Roche) three times for 45 s each at 4500 RPM with 2 min rests at 4°C. ∼300 µL of supernatant was moved into a clean tube, 300 µL of 70% ethanol was added, and samples were processed using the RNeasy Plus Mini Kit (QIAGEN) according to the manufacturer’s instructions. Sequencing libraries were prepared at the Tufts University Core Facility (TUCF Genomics) using the TruSeq Stranded mRNA Library Prep Kit (Illumina). 50-bp single-end reads were sequenced on an Illumina HiSeq 2500.

#### RNA-seq analysis

RNA-seq data analyses were performed using the Snakemake workflow management system (Koster and Rahmann, 2012), with code available on GitHub (github.com/khalillab). Additionally, an archive containing code and raw data suitable for reproducing all ChIP- and RNA-seq analyses is being submitted to Zenodo.

#### RNA-seq library processing

Adapter removal and 3’ quality trimming were performed using cutadapt (http://journal.embnet.org/index.php/embnetjournal/article/view/200). Reads were aligned using TopHat2 without a reference transcriptome, against *S. cerevisiae* genome build R64-2-1 modified to include the Venus reporter at the URA3 locus. Uniquely mapping reads were selected using SAMtools (Li et al., 2009). Coverage of the 5’-most base of the read (3’-most base of the RNA fragment) was extracted using bedtools genomecov (Quinlan and Hall, 2010), and normalized to the total number of uniquely mapped reads. Quality statistics of raw, cleaned, non-aligning, and uniquely aligning reads were assessed using FastQC (https://www.bioinformatics.babraham.ac.uk/projects/fastqc/).

#### RNA-seq differential expression analysis

RNA-seq differential expression analysis was performed for transcripts of verified coding genes, using an annotation of transcript boundaries based on TIF- (Pelechano et al., 2013) and TSS-seq (Doris et al., 2018) data that was modified to accommodate the Venus reporter at URA3. Read counts over these transcripts were input to differential expression analysis with DESeq2 (Love et al., 2014).

### Thermodynamic model

#### Model description

We constructed a thermodynamic model to gain insight into how cooperative assembly could be used to engineer specific regulatory connections in gene circuits, drawing on the rich history of describing transcriptional regulation by a thermodynamic treatment (Bintu et al., 2005; Buchler et al., 2003; Gertz et al., 2009; Shea and Ackers, 1985). For this study, we adapted our previously described model framework for cooperatively interacting synthetic transcription factors (synTFs) in yeast (Bashor et al., 2019).

Our model is composed of four key parameters: transcription factor concentration ([TF]), TF-DNA affinity (*K*_TF_), TF-TF cooperativity (*c*), and the number of binding sites at a given locus (*n*). We begin by enumerating all possible TF-bound promoter configurations for *n* binding sites. Each promoter state is assigned a transcriptional rate (*r*) and a thermodynamic weight (*w*). The transcriptional rate for a particular state is proportional to the number of TFs bound to a promoter. For simplicity, maximum transcriptional rate for a promoter is set to 1. Transcriptional weights describe the relative free energies of each state and are computed based on the number and affinity of interactions within each state, as previously described (Bashor et al., 2019). Transcriptional output is computed by averaging the relative contributions from each TF-bound promoter state:

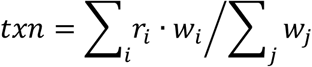

where *i* are transcriptionally active states and *j* are all promoter states.

To model cooperative synTF assembly, we include a promoter state weighted by an additional cooperativity (*c*) term. In our study, this could represent the additional free energy contribution by the clamp molecule on fully bound promoters. We assume that cooperativity does not affect the maximum rate of transcription at a promoter. Total transcription is calculated as described above. An example calculation of transcription at *n*=1 binding site is shown in **Figure S2A** and transcription at *n*=4 binding sites with cooperativity is shown in **Figure S2B**.

#### Regulatory specificity

We extended our model to investigate how biophysical parameters governing synTF assembly could be used to design specific regulatory connections. The model considers the simplified case of a TF that can interact with binding sites at a target synthetic (SYN) locus and an ‘off-target’ native (NAT) locus (**Figures 2A and S2**). To model different synTF assembly sizes, we considered a SYN locus with binding site clusters of *n =* 3 – 5; to model the spurious appearance of a CRM in the genome, we considered a NAT locus with *n =* 1 binding site.

We defined a regulatory specificity score as the difference between transcriptional output at the SYN (*txn_SYN_*) and NAT (*txn_NAT_*) loci (**Figure S2C**). Using the thermodynamic model, we computed regulatory specificity scores across a range of DNA affinities (*K*_TF_ = 10^−2^ – 10^2^ µM), cooperativities (*c* = 0 – 20 *k_B_T*), and SYN binding site numbers (*n* = 3 – 5). For simplicity, TF concentration was set to 1 µM for all simulations. We repeated this analysis for different formulations of the regulatory specificity score (**Figure S2D**).

All MATLAB code associated with this model is available on GitHub (github.com/khalillab/coop-TF-specificity).

### Population genetics model

We developed a population genetics model to explain the observed fitness dynamics in the eVOLVER continuous culture experiments. Generally, the dynamics of mutant progenies in adapting populations are shaped by both deterministic (e.g., natural selection) and stochastic forces (e.g., demographic fluctuations). It can be shown that the dynamics of a mutant progeny will be dominated by fluctuations when the population size is less than the inverse selective advantage (defined as the normalized fitness difference between the mutant and functional population) (Desai et al., 2007). In our experiments, a new mutant cell will obtain a fitness advantage on a time-scale comparable to the doubling time. As a result, the mutant population will grow deterministically after about one doubling and we can safely neglect demographic fluctuations (that is, fluctuations caused by finite cell numbers). To this end, we use an Ordinary Differential Equation (ODE) model to describe the population genetics.

In our model, we assume that cells grow at a rate *F*(*z*) where *z* is the concentration of a synTF. Before a synTF is induced, cells double approximately every 1.5 hrs (*λ*). After induction, cells pay a fitness penalty proportional to the fraction of ‘off-target’ NAT sites that are occupied by synTFs. We model off-target binding using our thermodynamic framework for *n* = 1 binding site, as before. This leads to the fitness function:

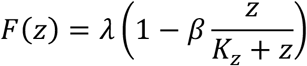

where *β* is the maximum fitness cost imposed by a synTF.

In our thermodynamic framework (**Figure S2**), *K*_TF_ can range from 10^−2^ to 10^2^ *μM*, but we work in units of TFs per cell. If a typical yeast cell is about 10^−15^ L, this translates to a range of 10^0^ to 10^4^ for *K*_*z*_. When induced, functional cells produce synTF at a rate *α* (per cell). Cells can mutate the transcriptional circuit at rate *μ* per unit time. We assume that the mutation rate does not depend on the doubling time. We know that the mutation rate per generation is roughly 3.5×10^−10^ (Lang and Murray, 2008). If we assume that there are ∼100s of potential mutants that can break the circuit, then the per hour rate to get a circuit-breaking mutation in a single lineage is 2.53×10^−8^.

Since there are roughly 10^8^ cells in the population, the average time to see a mutation is on the order of 1 h. Since cells are grown in our continuous culture experiments for 18 h prior to induction, it is reasonable to expect mutants in the population at the time of induction. However, prior to induction the mutations are nearly neutral (they incur no fitness benefit) and therefore the size of the mutant lineages will be determined solely by stochastic fluctuations. Standard theory dictates that if a mutant colony survives until the time of induction, its size will be on the order of the number of generations between the mutation and induction. This will be on the order of 100 cells, which we take as the initial mutant clone size. Using this order-of-magnitude estimate will be sufficient for our purposes, since we are ultimately interested in predicting qualitative features of the dynamics.

We model the number of functional (*x_f_*) and mutant (*x_m_*) cells in a growing population as:

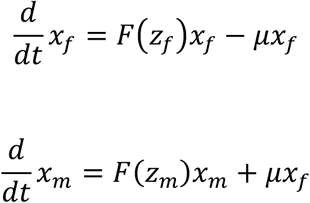

Letting TF_*f*_ denote the number of synTFs in functional cells, we have

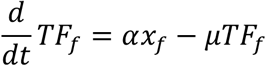

The second term comes from the fact that *μx*_*f*_ functional cells mutate per unit time, each taking *z*_*f*_=TF_*f*_ / *x*_*f*_ synTFs with it. Therefore, the number of synTFs in functional cells is described by the following equation:

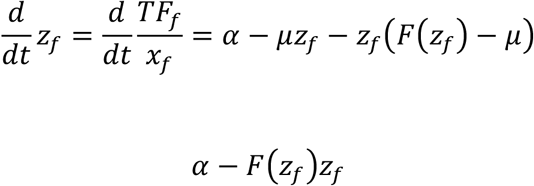

The number of synTFs in mutant (broken-circuit) cells is defined by:

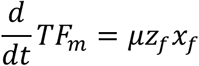

This implies that the change in the number of mutant (broken-circuit) cells can be described by:

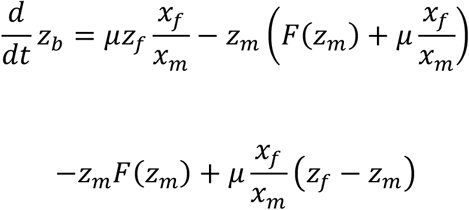

Note that we have assumed the population is growing exponentially, rather than in a finite culture. However, from the perspective of TF concentration and fitness only the species fractions, which are identical for exponentially growing and finite populations, are relevant.

We tested the qualitative features of this model by varying the maximum fitness cost (*β*) and synTF binding affinity (*K*_*z*_) while holding all other parameters constant. Changing the maximum fitness cost (**Figure S6F**) is analogous to choosing a different member of our zinc finger library, with varying DNA-binding specificities that could lead to differing levels of ‘OFF’-target interactions. Strains with a maximum high fitness cost show a severe growth defect after induction and are quickly out competed. This simulation result is similar to our eVOLVER continuous culture experiment using a second, high affinity zinc finger (13-6) (**Figure S6C**). Strains with comparably lower fitness costs are also lost over time but at a slower rate. Changing synTF binding affinity in the model is analogous to testing the high and low affinity ZF variants. As with the evolution experiments, functional cells harboring low affinity synTFs are retained for longer periods of time compared to those with high affinity synTFs.

## QUANTIFICATION AND STATISTICAL ANALYSIS

FlowJo was used to extract geometric mean fluorescence values or the percentage of mTurquoise, mVenus or mRuby activated cells from flow cytometry measurements. Microsoft Excel and GraphPad Prism software were used to process data. Statistical details such as number of replicates and error calculations are provided in figure legends.

## DATA AND SOFTWARE AVAILABILITY

Raw RNA-seq and ChIP-seq data for transcriptome and binding analyses, respectively, are accessible at NCBI GEO database, accession GSE203146.

## SUPPLEMENTAL DATA

**Supplemental Table S1. Related to Figures 1-6.** Plasmids used in this study.

**Supplemental Table S2. Related to Figures 1-6.** Yeast strains used in this study.

## SUPPLEMENTAL FIGURES

**Figure S1.**
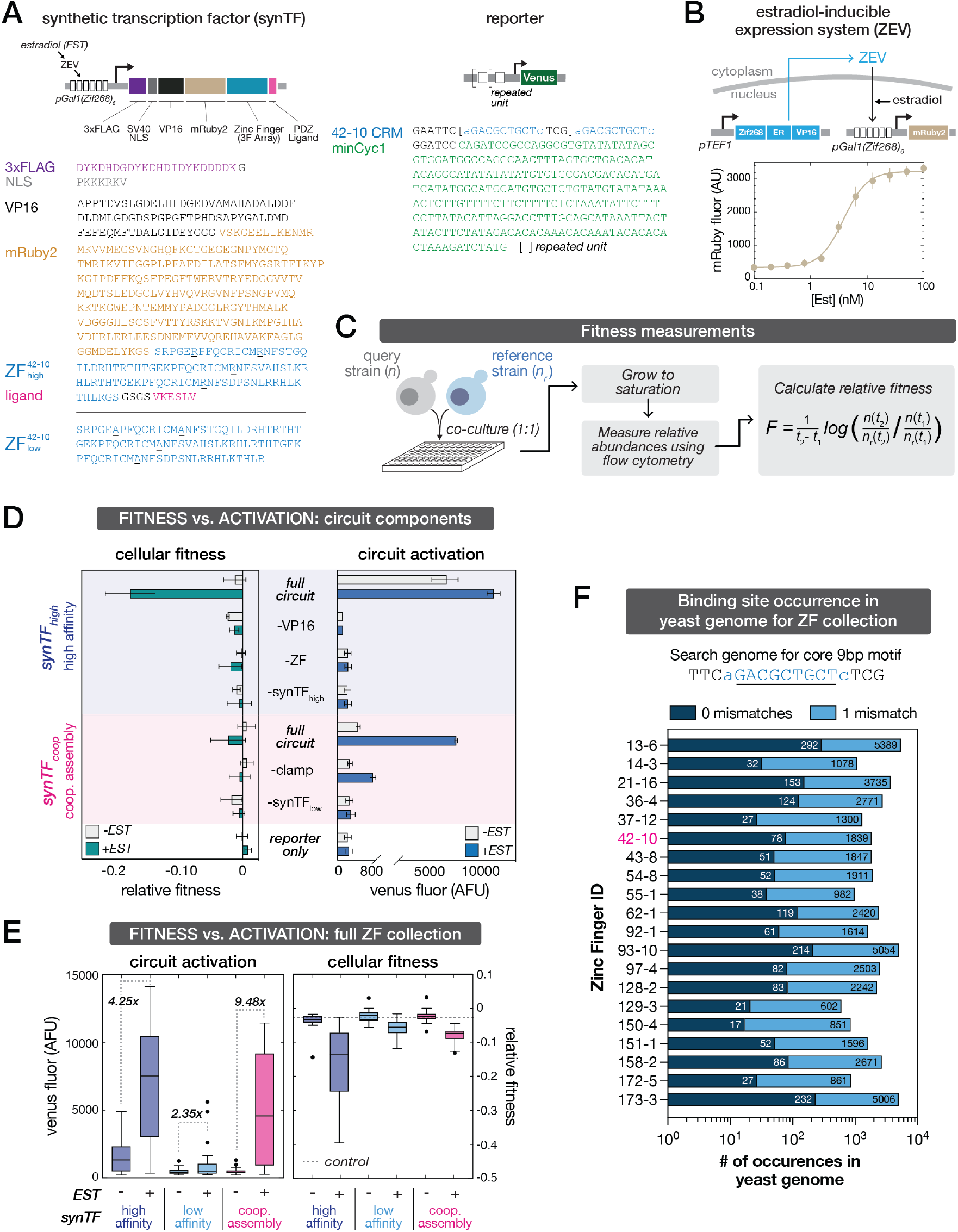
Design and characterization of synthetic circuits. (A) Design details of inducible synTF circuit components including amino acid sequences common to all synTFs used in this study. SynTF activators are composed of a 3-finger (3F) ZF array fused to a herpes simplex VP16 transactivation domain (TAD), 3x repeat FLAG epitope tag (Sigma), mRuby fluorescent protein, syntrophin PDZ ligand, and SV40-derived nuclear localization sequence (NLS). For zinc finger affinity alleles, mutated arginine residues are underlined. DNA sequences for engineered synTF promoters, composed of a minimal CYC1 (minCyc1) promoter with upstream synTF binding sequences (CRMs). All sequences were yeast-optimized and chromosomally integrated into *S. cerevisiae*. (B) Characterization of inducible expression system. mRuby2 fluorescent protein expression was used as a proxy to quantify the ß-estradiol-inducible promoter. Flow cytometry measurements were made at mid-log phase and error bars indicate standard deviation from three biological replicates. [Est], ß-estradiol concentration. (C) Workflow for the competition co-culture experiment used to quantify fitness across this study. Query strains (and associated control strains) were each co-cultured 1:1 with a reference strain (parental strain constitutively expressing mTurquoise reporter). Three biological replicates (separate colonies) of each query or control strain were measured in separate wells, and duplicate experiments were performed in media with and without 1 uM ß-estradiol inducer. Cocultures were sampled when initially mixed (T0) and every 12 hours to determine relative abundancies of reference vs. query or control strain, derived from the fraction of cells in each well expressing mTurquoise. Fitness measurements were equated for each query strain by calculating changes in mTurquoise expression from T0 to the experimental endpoint. (D) Deletion of either the VP16 transactivation domain or the ZF DNA-binding domain is sufficient to rescue the fitness cost induced by synTF_high_ circuits. Strains lacking designated circuit components were constructed, and cellular fitness and circuit activation measured as previously described. Bars represent mean values for three biological replicates +/− SD (measured in two separate experiments). (E) Cellular fitness and circuit activation for synTF circuits constructed from a collection of distinct ZF species. Fitness and activation profiles were quantified at 36 hours following induction, in conditions with and without ß-estradiol inducer. In this case, ‘control’ denotes a strain with no integrated circuit components, since reporter-only strains are different for each synTF. Tukey boxplots represent the range of means for 20 synTF ZF library members. (F) Prevalence of synthetic zinc finger (ZF) binding sites in the yeast genome. Occurrences of the full and single mismatches of the predicted core (9-bp) binding motifs for the full ZF collection.

**Figure S2.**
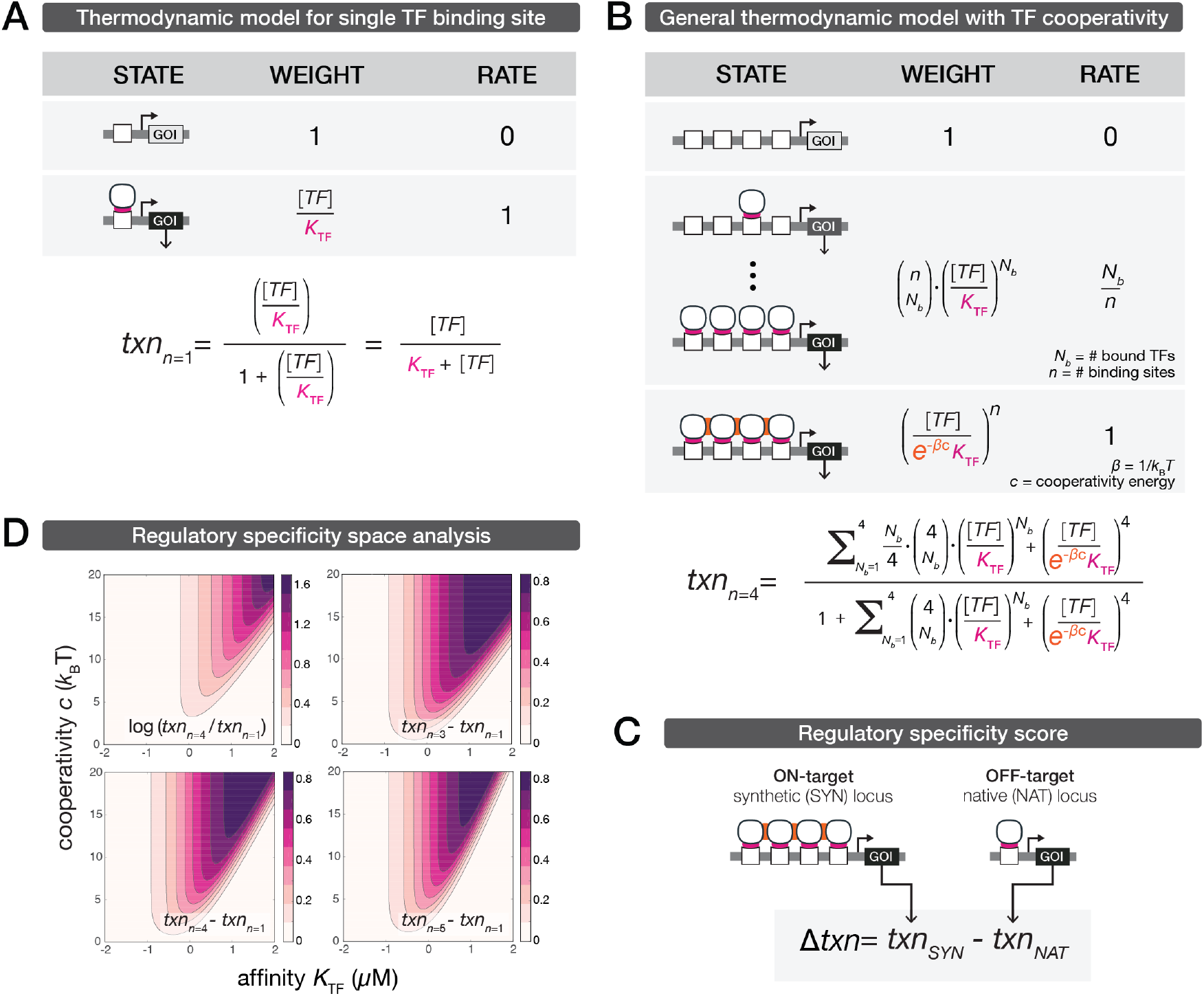
Description and analysis of thermodynamic model for cooperative assembly-driven specificity. (A) Description of thermodynamic model for a single TF binding site. Grey boxes: Enumeration of promoter states with corresponding thermodynamic weights and transcriptional rates (proportional to the number of bound TFs). Equation: Transcriptional output (*txn*_n=1_) is obtained by averaging the relative transcriptional contributions of all promoter states. *K*_TF_, TF-DNA binding affinity. (B) Description of a generalized thermodynamic model incorporating TF cooperative assembly. Grey boxes: Enumeration of promoter states with corresponding thermodynamic weights and transcriptional rates (proportional to the number of bound TFs). Equation: Transcriptional output for *n*=4 binding sites. *K*_TF_, TF-DNA binding affinity; *c*, cooperativity term that defines the additional stability provided by the multivalent TF interactions. (C) Regulatory specificity score is defined as the difference between SYN on- vs. NAT off-target transcription. (D) Regulatory specificity space is qualitatively preserved for different model formulations. Alternative regulatory specificity score with *n*=4 binding sites (top left). Specificity score as described in Fig. S2C for different number of binding sites at the SYN locus: *n*=3 (top right), *n*=4 (bottom left), and *n*=5 (bottom right).

**Figure S3.**
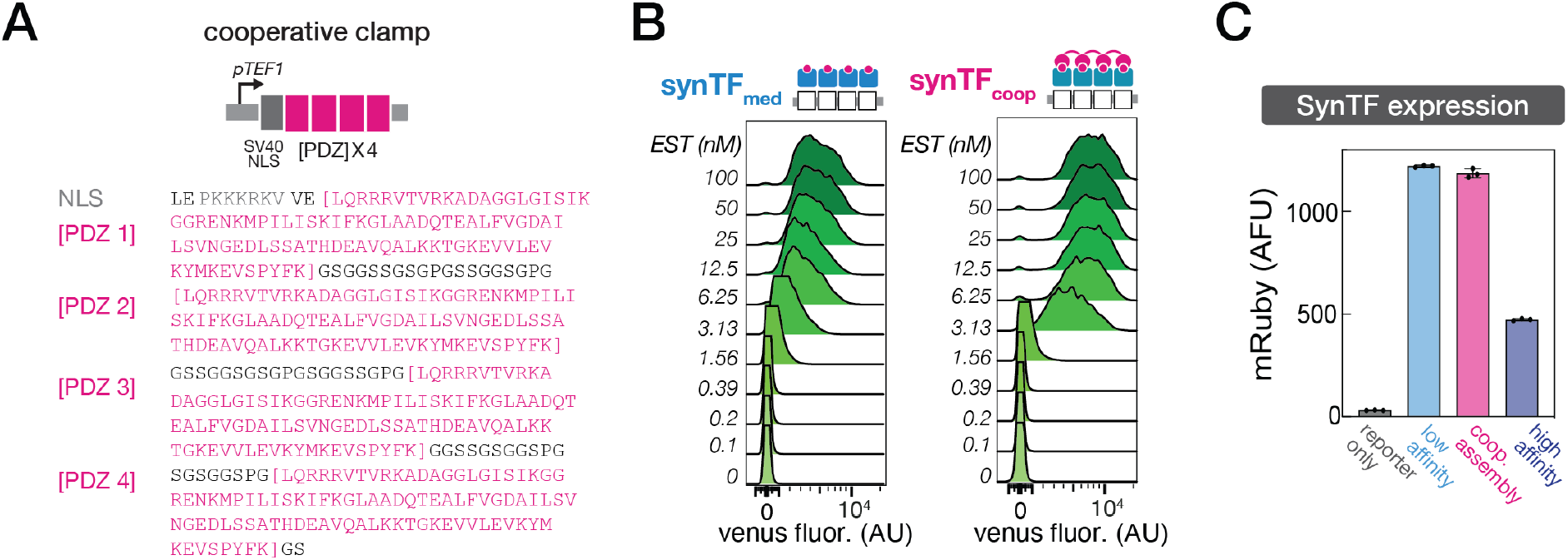
Design of the cooperative clamp and characterization of clamp-mediated cooperative synthetic circuits. (A) Design and sequence details of the cooperative clamp. Cooperative clamps are composed of the same SV40-derived NLS followed by repeat units of the syntrophin PDZ domain. An *n* = 4 clamp sequence is depicted with the repeat units highlighted. Sequences were yeast-optimized and chromosomally integrated into *S. cerevisiae*. (B) Single-cell dose response behaviors for the independent (synTF_med_) and cooperative (synTF_coop_) synTF circuits. Flow cytometry analysis of the dose responses show a linear shift from OFF to ON with the non-clamp synTF and a characteristically non-linear shift from OFF to ON with the clamp-mediated cooperative synTF architecture in response to increasing inducer concentrations. (C) SynTF expression levels measured by quantifying synTF-mRuby2 fluorescence following circuit induction. Bars represent mean values for three biological replicates +/− SD.

**Figure S4.**
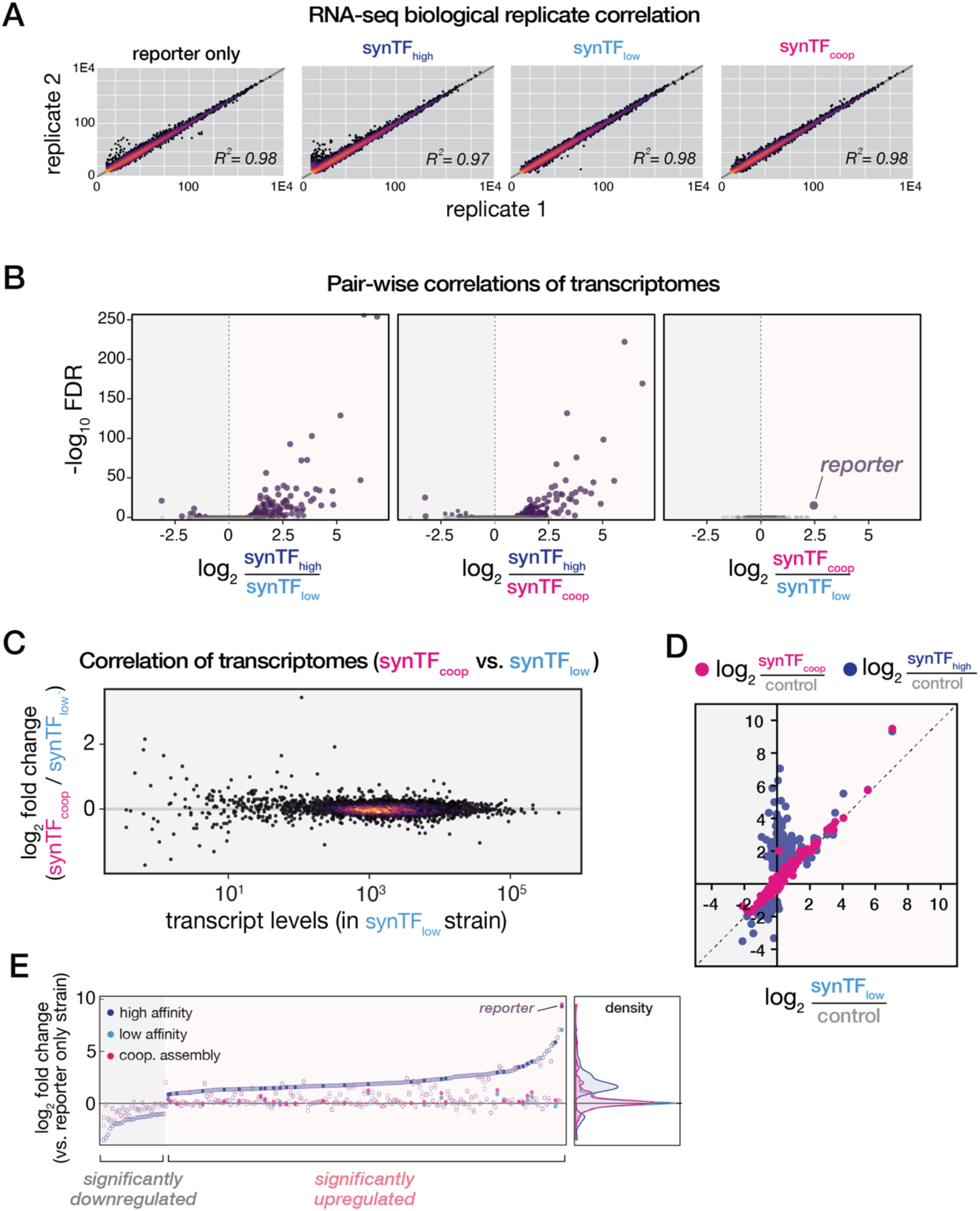
Correlation of transcriptomes of strains expressing synTF circuit variants. (A) Correlation of RNA-seq measurements for biological replicates. Correlation coefficients were calculated using log_2_ transformed expression values. (B) Pair-wise correlations of transcriptomes for the three synTF circuit configurations. The synthetic reporter was the only gene differentially regulated by synTF_coop_ vs. synTF_low_. Two biological replicates for each strain are reported. FDR, false discovery rate. (C) Transcript levels are highly correlated between strains expressing synTF_coop_ and synTF_low_ circuits. (D) Differential gene expression values for synTF_high_ and synTF_coop_ against synTF_low_. Differential gene expression for the synTF_coop_ strain correlates highly with synTF_low_, with one notable exception: the synthetic reporter gene, which is differentially expressed to equivalent levels as the synTF_high_ strain. (E) Differential gene expression analysis of RNA-seq measurements following induction of synTF_high_, synTF_low_, and synTF_coop_ circuits. Plotted are genes that are significantly differentially regulated relative to the reporter-only control. The synTF_high_ circuit induces a global misregulation of the host transcriptome, including significantly upregulating 182 genes. Gene expression density distributions of synTF_coop_ and synTF_low_ strains are highly similar to one another and cluster tightly around the reporter-only background. Filled circles represent genes with a motif located within 300-bp upstream of the TSS that has at least 8/9 bp homology to the cognate motif.

**Figure S5.**
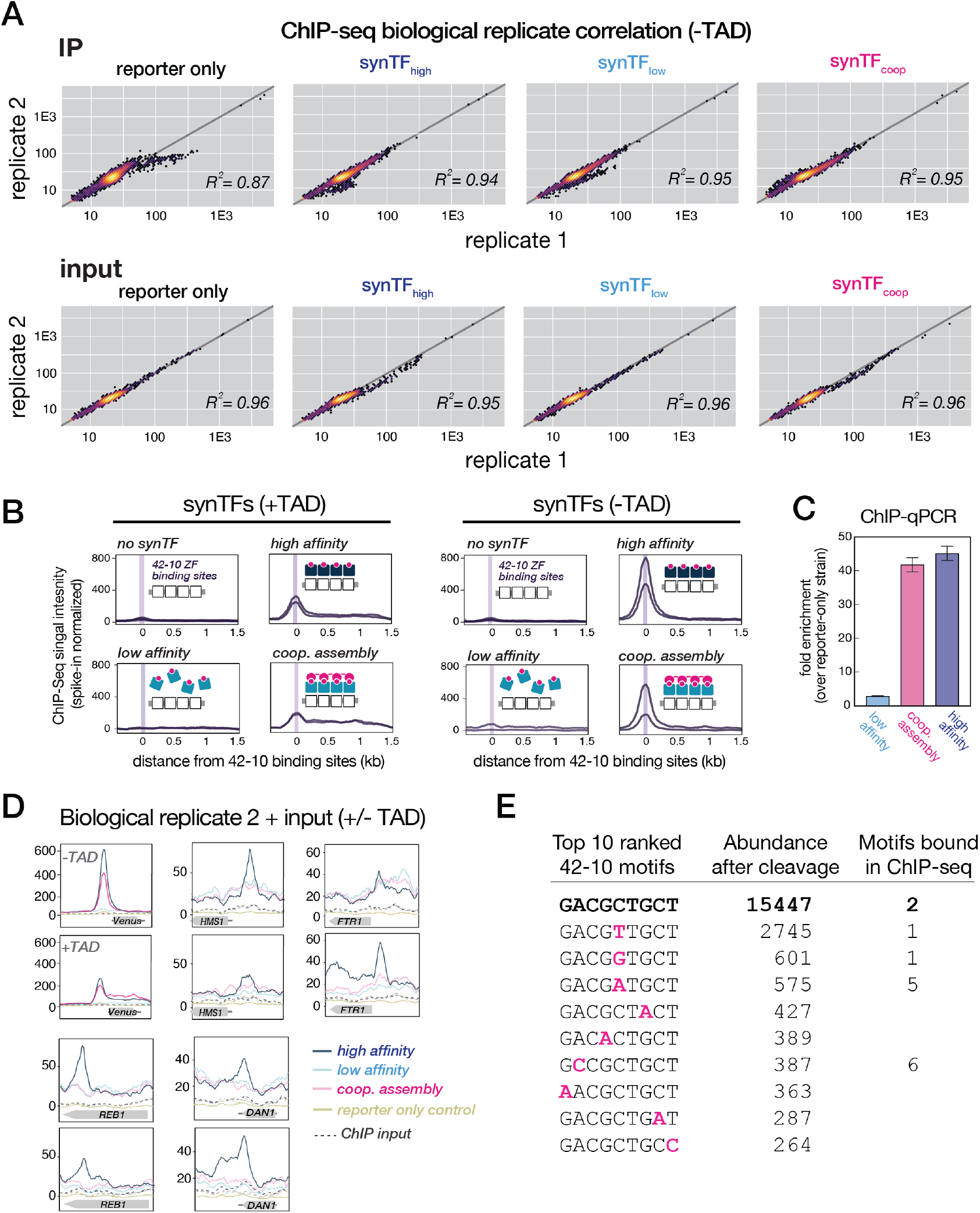
Validation of ChIP-seq methods and synTF binding profiles. (A) Correlation of ChIP-seq measurements for biological replicates. Correlations are plotted for both immunoprecipitated (IP) and input control samples for synTFs lacking a TAD fusion. Correlation coefficients were calculated using log_2_ transformed expression values. (B) ChIP enrichment profiles at the synthetic reporter locus for no synTF, synTF_low_, synTF_high_, and synTF_coop_ (-/+ TAD). Binding enrichment for two biological replicates of each strain are shown. The maximum peak heights (purple line) are highly correlated with the genomic location of the ZF 42-10 binding sites. Relative ChIP enrichment was normalized to FLAG-tagged *S. pombe* spike-in DNA that was produced in parallel with the *S. cerevisiae* samples. (C) SynTF enrichment at the synthetic reporter locus measured using ChIP-quantitative PCR (ChIP-qPCR). Fold enrichment is determined for each condition compared to a reporter-only control. Enrichment patterns recapitulate those observed with ChIP-seq. Bars represent mean values for three technical replicates +/− SD. (D) ChIP enrichment profiles for each synTF (low, high, coop) strain, reporter-only control strain, and input control sample at five representative loci. Relative ChIP enrichment was normalized to FLAG-tagged *S. pombe* spike-in DNA that was produced in parallel with the *S. cerevisiae* samples. (E) Top-ranked interaction motifs for ZF 42-10, as determined by an independent dataset based on an *in vitro* DNA cleavage profiling assay (Pattanayak et al., 2011). Abundance after cleavage quantifies the frequency that a motif has been targeted (and cleaved) by a nuclease version of our candidate ZF. The 15 synTF_high_ binding events nominated by our ChIP-seq analysis occurred at the seven most preferred motifs.

**Figure S6.**
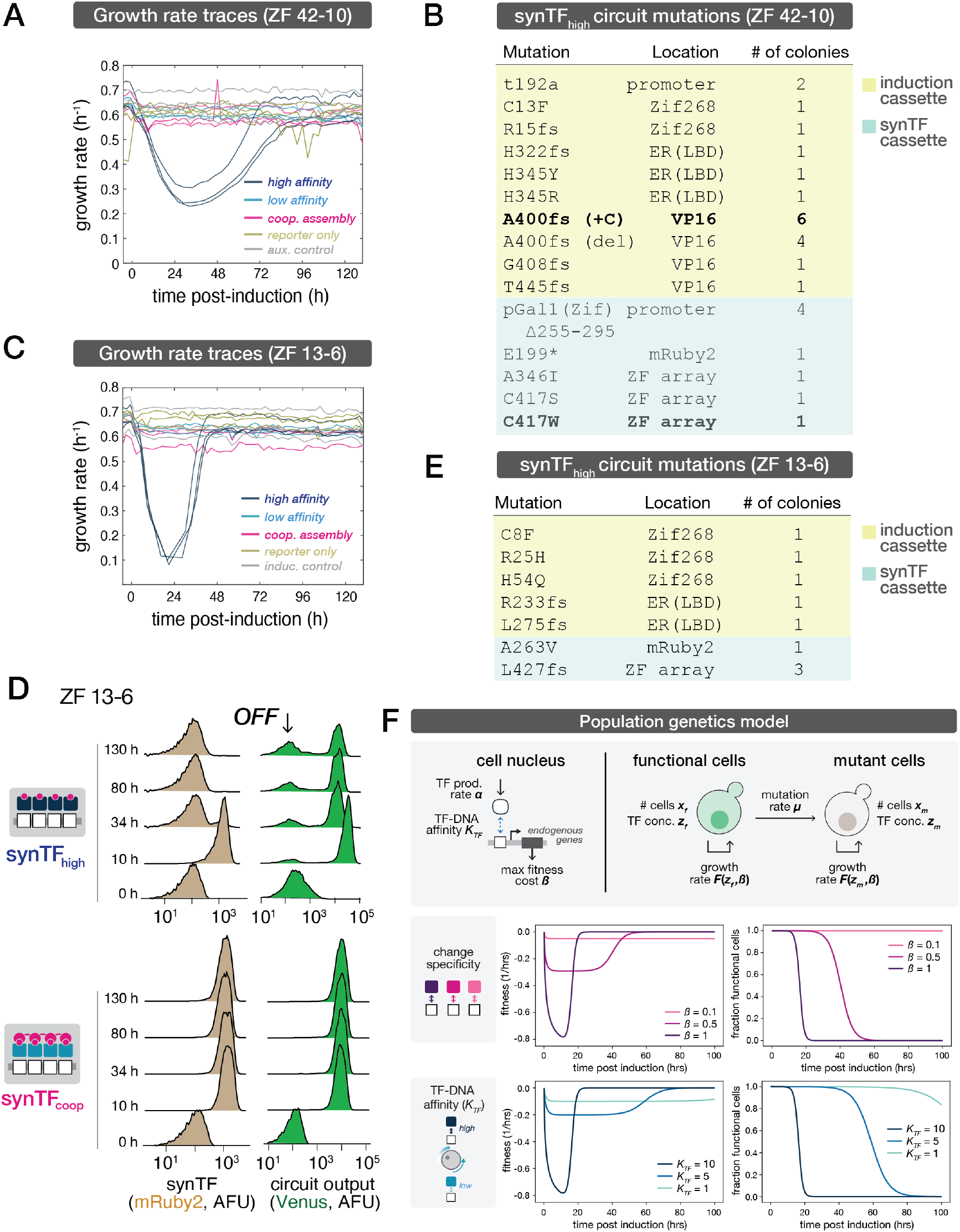
Growth, circuit expression, and mutational patterns are conserved in a replicate long-term culture experiment and captured by a population genetics model. (A) Raw growth rate traces for three biological replicates of synTF and control strains in long-term eVOLVER continuous culture. Each replicate was cultured in a separate eVOLVER vial. The auxiliary (aux.) control has a scrambled placeholder sequence integrated into each of the three loci into which circuit components are integrated. (B) Mutational analysis of the circuit genotype from synTF_high_-derived colonies following eVOLVER long-term culture. Mutations were identified within the induction cassette (yellow) and synTF cassette (green). A single residue in the induction cassette was highly targeted, with mutations identified in 10 of the sequenced colonies. (C) Raw growth rate traces for three biological replicates of a second synTF species (ZF 13-6) and control strains in long-term eVOLVER continuous culture. Each replicate was cultured in a separate eVOLVER vial. The inducer (induc.) control has the induction cassette driving an mRuby2 fluorescent reporter in place of the same induction cassette driving the synTFs. (D) Single-cell flow cytometry distributions of synTF and circuit reporter expression over the time course of the continuous culture experiment for a second synTF species (ZF 13-6). (E) Mutational analysis of the circuit genotype from synTF_high_-derived colonies following eVOLVER long-term culture with a second synTF species (ZF 13-6). (F) A population genetics model captures the population fitness and circuit retention dynamics observed in long-term culture experiments. Description of the model (top). A TF is produced at a constant rate (*α*) and has an affinity for DNA (*K_TF_*), which is proportional to the maximum fitness cost (*β*) it imposes on a host cell. The number of functional and mutant cells in a population is defined by *x_f_* and *x_m_*, respectively. The concentration of TF in each cell type is defined by *z_f_* and *z_m_*, respectively. Functional cells are converted to mutant cells at a constant rate *µ*. The growth rate of each population (*F*) is a function of the concentration and maximum fitness cost of each TF. Population fitness (middle left) and circuit retention (middle right) dynamics for a range of TF fitness cost values (*β*), where *K_TF_* = 1. Population fitness (bottom left) and circuit retention (bottom right) dynamics for a range of TF-DNA binding affinities (*K_TF_*), where *β* = 1.

