## Supplementary material for "Cooperative assembly confers regulatory specificity and long-term genetic circuit stability": Table S1

**Table S1. Related to Figures 1-6.** Plasmids used in this study

| Plasmid | Content | Bacterial selection | Yeast selection | Yeast plasmid type | Integration locus | Restriction site (integration) | Purpose | Reference |
|---|---|---|---|---|---|---|---|---|
| pKLE-01 | empty | Amp | -ura | Integrating | URA3 | NotI | Parent plasmid for single-integration at URA3 | This study |
| pKLE-09 | empty | Amp | hygromycinR | Integrating | HO | NotI | Parent plasmid for single-integration at HO (for pKL1s | This study |
| pKLE-15 | empty | Kan | hygromycinR | Integrating | HO | NotI | Parent plasmid for single-integration at HO (for pKL2s | This study |
| pKLE-22 | empty | Amp | -leu | Integrating | LEU2 | NotI | Parent plasmid for single-integration at LEU2 | This study |
| pKL1-010 | {pTDH3-mTurquoise2-tSSA1} | Amp | hygromycinR | Integrating | HO | NotI | Constitutive mTurquoise reference strain for fitness ex | This study |
| pKL1-065 | {Spacer} | Amp | -ura | Integrating | URA3 | NotI | Spacer sequence for ura locus | This study |
| pKL1-095 | {(13-6 op)x4-minCyc1-Venus-tCyc1} | Amp | -ura | Integrating | URA3 | NotI | Venus fluorescent reporter with 4x CRMs | This study |
| pKL1-096 | {(14-3 op)x4-minCyc1-Venus-tCyc1} | Amp | -ura | Integrating | URA3 | NotI | Venus fluorescent reporter with 4x CRMs | This study |
| pKL1-097 | {(21-16 op)x4-minCyc1-Venus-tCyc1} | Amp | -ura | Integrating | URA3 | NotI | Venus fluorescent reporter with 4x CRMs | This study |
| pKL1-098 | {(36-4 op)x4-minCyc1-Venus-tCyc1} | Amp | -ura | Integrating | URA3 | NotI | Venus fluorescent reporter with 4x CRMs | This study |
| pKL1-099 | {(37-12 op)x4-minCyc1-Venus-tCyc1} | Amp | -ura | Integrating | URA3 | NotI | Venus fluorescent reporter with 4x CRMs | This study |
| pKL1-100 | {(42-10 op)x4-minCyc1-Venus-tCyc1} | Amp | -ura | Integrating | URA3 | NotI | Venus fluorescent reporter with 4x CRMs | This study |
| pKL1-101 | {(43-8 op)x4-minCyc1-Venus-tCyc1} | Amp | -ura | Integrating | URA3 | NotI | Venus fluorescent reporter with 4x CRMs | This study |
| pKL1-102 | {(54-8 op)x4-minCyc1-Venus-tCyc1} | Amp | -ura | Integrating | URA3 | NotI | Venus fluorescent reporter with 4x CRMs | This study |
| pKL1-103 | {(55-1 op)x4-minCyc1-Venus-tCyc1} | Amp | -ura | Integrating | URA3 | NotI | Venus fluorescent reporter with 4x CRMs | This study |
| pKL1-104 | {(62-1 op)x4-minCyc1-Venus-tCyc1} | Amp | -ura | Integrating | URA3 | NotI | Venus fluorescent reporter with 4x CRMs | This study |
| pKL1-105 | {(63-4 op)x4-minCyc1-Venus-tCyc1} | Amp | -ura | Integrating | URA3 | NotI | Venus fluorescent reporter with 4x CRMs | This study |
| pKL1-106 | {(92-1 op)x4-minCyc1-Venus-tCyc1} | Amp | -ura | Integrating | URA3 | NotI | Venus fluorescent reporter with 4x CRMs | This study |
| pKL1-107 | {(93-10 op)x4-minCyc1-Venus-tCyc1} | Amp | -ura | Integrating | URA3 | NotI | Venus fluorescent reporter with 4x CRMs | This study |
| pKL1-108 | {(97-4 op)x4-minCyc1-Venus-tCyc1} | Amp | -ura | Integrating | URA3 | NotI | Venus fluorescent reporter with 4x CRMs | This study |
| pKL1-109 | {(128-2 op)x4-minCyc1-Venus-tCyc1} | Amp | -ura | Integrating | URA3 | NotI | Venus fluorescent reporter with 4x CRMs | This study |
| pKL1-110 | {(129-3 op)x4-minCyc1-Venus-tCyc1} | Amp | -ura | Integrating | URA3 | NotI | Venus fluorescent reporter with 4x CRMs | This study |
| pKL1-111 | {(150-4 op)x4-minCyc1-Venus-tCyc1} | Amp | -ura | Integrating | URA3 | NotI | Venus fluorescent reporter with 4x CRMs | This study |
| pKL1-112 | {(151-1 op)x4-minCyc1-Venus-tCyc1} | Amp | -ura | Integrating | URA3 | NotI | Venus fluorescent reporter with 4x CRMs | This study |
| pKL1-113 | {(158-2 op)x4-minCyc1-Venus-tCyc1} | Amp | -ura | Integrating | URA3 | NotI | Venus fluorescent reporter with 4x CRMs | This study |
| pKL1-116 | {(173-3 op)x4-minCyc1-Venus-tCyc1} | Amp | -ura | Integrating | URA3 | NotI | Venus fluorescent reporter with 4x CRMs | This study |
| pKL2-133 | {pTEF1-ZEV}{pGal1(Zif268op)x6-3xFLAG-NLS-VP16-mRuby2-13-6-4x-PDZLigS5*-tSSA1} | Kan | hygromycinR | Integrating | HO | NotI | 4x R->A mutant ZF synTF | This study |
| pKL2-134 | {pTEF1-ZEV}{pGal1(Zif268op)x6-3xFLAG-NLS-VP16-mRuby2-14-3-4x-PDZLigS5*-tSSA1} | Kan | hygromycinR | Integrating | HO | NotI | 4x R->A mutant ZF synTF | This study |
| pKL2-135 | {pTEF1-ZEV}{pGal1(Zif268op)x6-3xFLAG-NLS-VP16-mRuby2-21-16-4x-PDZLigS5*-tSSA1} | Kan | hygromycinR | Integrating | HO | NotI | 4x R->A mutant ZF synTF | This study |
| pKL2-136 | {pTEF1-ZEV}{pGal1(Zif268op)x6-3xFLAG-NLS-VP16-mRuby2-36-4-4x-PDZLigS5*-tSSA1} | Kan | hygromycinR | Integrating | HO | NotI | 4x R->A mutant ZF synTF | This study |
| pKL2-137 | {pTEF1-ZEV}{pGal1(Zif268op)x6-3xFLAG-NLS-VP16-mRuby2-37-12-4x-PDZLigS5*-tSSA1} | Kan | hygromycinR | Integrating | HO | NotI | 4x R->A mutant ZF synTF | This study |
| pKL2-138 | {pTEF1-ZEV}{pGal1(Zif268op)x6-3xFLAG-NLS-VP16-mRuby2-42-10-4x-PDZLigS5*-tSSA1} | Kan | hygromycinR | Integrating | HO | NotI | 4x R->A mutant ZF synTF | This study |
| pKL2-139 | {pTEF1-ZEV}{pGal1(Zif268op)x6-3xFLAG-NLS-VP16-mRuby2-43-8-4x-PDZLigS5*-tSSA1} | Kan | hygromycinR | Integrating | HO | NotI | 4x R->A mutant ZF synTF | This study |
| pKL2-140 | {pTEF1-ZEV}{pGal1(Zif268op)x6-3xFLAG-NLS-VP16-mRuby2-54-8-4x-PDZLigS5*-tSSA1} | Kan | hygromycinR | Integrating | HO | NotI | 4x R->A mutant ZF synTF | This study |
| pKL2-141 | {pTEF1-ZEV}{pGal1(Zif268op)x6-3xFLAG-NLS-VP16-mRuby2-55-1-4x-PDZLigS5*-tSSA1} | Kan | hygromycinR | Integrating | HO | NotI | 4x R->A mutant ZF synTF | This study |
| pKL2-142 | {pTEF1-ZEV}{pGal1(Zif268op)x6-3xFLAG-NLS-VP16-mRuby2-62-1-4x-PDZLigS5*-tSSA1} | Kan | hygromycinR | Integrating | HO | NotI | 4x R->A mutant ZF synTF | This study |
| pKL2-143 | {pTEF1-ZEV}{pGal1(Zif268op)x6-3xFLAG-NLS-VP16-mRuby2-63-4-4x-PDZLigS5*-tSSA1} | Kan | hygromycinR | Integrating | HO | NotI | 4x R->A mutant ZF synTF | This study |
| pKL2-144 | {pTEF1-ZEV}{pGal1(Zif268op)x6-3xFLAG-NLS-VP16-mRuby2-92-1-4x-PDZLigS5*-tSSA1} | Kan | hygromycinR | Integrating | HO | NotI | 4x R->A mutant ZF synTF | This study |
| pKL2-145 | {pTEF1-ZEV}{pGal1(Zif268op)x6-3xFLAG-NLS-VP16-mRuby2-93-10-4x-PDZLigS5*-tSSA1} | Kan | hygromycinR | Integrating | HO | NotI | 4x R->A mutant ZF synTF | This study |
| pKL2-146 | {pTEF1-ZEV}{pGal1(Zif268op)x6-3xFLAG-NLS-VP16-mRuby2-97-4-4x-PDZLigS5*-tSSA1} | Kan | hygromycinR | Integrating | HO | NotI | 4x R->A mutant ZF synTF | This study |
| pKL2-147 | {pTEF1-ZEV}{pGal1(Zif268op)x6-3xFLAG-NLS-VP16-mRuby2-128-2-4x-PDZLigS5*-tSSA1} | Kan | hygromycinR | Integrating | HO | NotI | 4x R->A mutant ZF synTF | This study |
| pKL2-148 | {pTEF1-ZEV}{pGal1(Zif268op)x6-3xFLAG-NLS-VP16-mRuby2-129-3-4x-PDZLigS5*-tSSA1} | Kan | hygromycinR | Integrating | HO | NotI | 4x R->A mutant ZF synTF | This study |
| pKL2-149 | {pTEF1-ZEV}{pGal1(Zif268op)x6-3xFLAG-NLS-VP16-mRuby2-150-4-4x-PDZLigS5*-tSSA1} | Kan | hygromycinR | Integrating | HO | NotI | 4x R->A mutant ZF synTF | This study |
| pKL2-150 | {pTEF1-ZEV}{pGal1(Zif268op)x6-3xFLAG-NLS-VP16-mRuby2-151-1-4x-PDZLigS5*-tSSA1} | Kan | hygromycinR | Integrating | HO | NotI | 4x R->A mutant ZF synTF | This study |
| pKL2-151 | {pTEF1-ZEV}{pGal1(Zif268op)x6-3xFLAG-NLS-VP16-mRuby2-158-2-4x-PDZLigS5*-tSSA1} | Kan | hygromycinR | Integrating | HO | NotI | 4x R->A mutant ZF synTF | This study |
| pKL2-154 | {pTEF1-ZEV}{pGal1(Zif268op)x6-3xFLAG-NLS-VP16-mRuby2-173-3-4x-PDZLigS5*-tSSA1} | Kan | hygromycinR | Integrating | HO | NotI | 4x R->A mutant ZF synTF | This study |
| pKL2-155 | {pTEF1-ZEV}{pGal1(Zif268op)x6-3xFLAG-NLS-VP16-mRuby2-13-6-WT-PDZLigS5*-tSSA1} | Kan | hygromycinR | Integrating | HO | NotI | WT ZF synTF | This study |
| pKL2-156 | {pTEF1-ZEV}{pGal1(Zif268op)x6-3xFLAG-NLS-VP16-mRuby2-14-3-WT-PDZLigS5*-tSSA1} | Kan | hygromycinR | Integrating | HO | NotI | WT ZF synTF | This study |
| pKL2-157 | {pTEF1-ZEV}{pGal1(Zif268op)x6-3xFLAG-NLS-VP16-mRuby2-21-16-WT-PDZLigS5*-tSSA1} | Kan | hygromycinR | Integrating | HO | NotI | WT ZF synTF | This study |
| pKL2-158 | {pTEF1-ZEV}{pGal1(Zif268op)x6-3xFLAG-NLS-VP16-mRuby2-36-4-WT-PDZLigS5*-tSSA1} | Kan | hygromycinR | Integrating | HO | NotI | WT ZF synTF | This study |
| pKL2-159 | {pTEF1-ZEV}{pGal1(Zif268op)x6-3xFLAG-NLS-VP16-mRuby2-37-12-WT-PDZLigS5*-tSSA1} | Kan | hygromycinR | Integrating | HO | NotI | WT ZF synTF | This study |
| pKL2-160 | {pTEF1-ZEV}{pGal1(Zif268op)x6-3xFLAG-NLS-VP16-mRuby2-42-10-WT-PDZLigS5*-tSSA1} | Kan | hygromycinR | Integrating | HO | NotI | WT ZF synTF | This study |
| pKL2-161 | {pTEF1-ZEV}{pGal1(Zif268op)x6-3xFLAG-NLS-VP16-mRuby2-43-8-WT-PDZLigS5*-tSSA1} | Kan | hygromycinR | Integrating | HO | NotI | WT ZF synTF | This study |
| pKL2-162 | {pTEF1-ZEV}{pGal1(Zif268op)x6-3xFLAG-NLS-VP16-mRuby2-54-8-WT-PDZLigS5*-tSSA1} | Kan | hygromycinR | Integrating | HO | NotI | WT ZF synTF | This study |
| pKL2-163 | {pTEF1-ZEV}{pGal1(Zif268op)x6-3xFLAG-NLS-VP16-mRuby2-55-1-WT-PDZLigS5*-tSSA1} | Kan | hygromycinR | Integrating | HO | NotI | WT ZF synTF | This study |
| pKL2-164 | {pTEF1-ZEV}{pGal1(Zif268op)x6-3xFLAG-NLS-VP16-mRuby2-62-1-WT-PDZLigS5*-tSSA1} | Kan | hygromycinR | Integrating | HO | NotI | WT ZF synTF | This study |
| pKL2-165 | {pTEF1-ZEV}{pGal1(Zif268op)x6-3xFLAG-NLS-VP16-mRuby2-63-4-WT-PDZLigS5*-tSSA1} | Kan | hygromycinR | Integrating | HO | NotI | WT ZF synTF | This study |
| pKL2-166 | {pTEF1-ZEV}{pGal1(Zif268op)x6-3xFLAG-NLS-VP16-mRuby2-92-1-WT-PDZLigS5*-tSSA1} | Kan | hygromycinR | Integrating | HO | NotI | WT ZF synTF | This study |

| | | | | | | | | |
|---|---|---|---|---|---|---|---|---|
| pKL2-167 | {pTEF1-ZEV}{pGal1(Zif268op)x6-3xFLAG-NLS-VP16-mRuby2-93-10-WT-PDZLigS5*-tSSA1} | Kan | hygromycinR | Integrating | HO | NotI | WT ZF synTF | This study |
| pKL2-168 | {pTEF1-ZEV}{pGal1(Zif268op)x6-3xFLAG-NLS-VP16-mRuby2-97-4-WT-PDZLigS5*-tSSA1} | Kan | hygromycinR | Integrating | HO | NotI | WT ZF synTF | This study |
| pKL2-169 | {pTEF1-ZEV}{pGal1(Zif268op)x6-3xFLAG-NLS-VP16-mRuby2-128-2-WT-PDZLigS5*-tSSA1} | Kan | hygromycinR | Integrating | HO | NotI | WT ZF synTF | This study |
| pKL2-170 | {pTEF1-ZEV}{pGal1(Zif268op)x6-3xFLAG-NLS-VP16-mRuby2-129-3-WT-PDZLigS5*-tSSA1} | Kan | hygromycinR | Integrating | HO | NotI | WT ZF synTF | This study |
| pKL2-171 | {pTEF1-ZEV}{pGal1(Zif268op)x6-3xFLAG-NLS-VP16-mRuby2-150-4-WT-PDZLigS5*-tSSA1} | Kan | hygromycinR | Integrating | HO | NotI | WT ZF synTF | This study |
| pKL2-172 | {pTEF1-ZEV}{pGal1(Zif268op)x6-3xFLAG-NLS-VP16-mRuby2-151-1-WT-PDZLigS5*-tSSA1} | Kan | hygromycinR | Integrating | HO | NotI | WT ZF synTF | This study |
| pKL2-173 | {pTEF1-ZEV}{pGal1(Zif268op)x6-3xFLAG-NLS-VP16-mRuby2-158-2-WT-PDZLigS5*-tSSA1} | Kan | hygromycinR | Integrating | HO | NotI | WT ZF synTF | This study |
| pKL2-176 | {pTEF1-ZEV}{pGal1(Zif268op)x6-3xFLAG-NLS-VP16-mRuby2-173-3-WT-PDZLigS5*-tSSA1} | Kan | hygromycinR | Integrating | HO | NotI | WT ZF synTF | This study |
| pKL1-222 | {pTEF1-NLS-(Syn)2-20GS-(Syn)2-tADH1} | Amp | -leu | Integrating | LEU2 | NotI | 4 x clamp | Bashor et al., 2019 & this stu |
| pKL1-224 | {Spacer} | Amp | -leu | Integrating | LEU2 | NotI | Spacer sequence for leu locus | This study |
| pKL2-020 | {pTEF1-ZEV}{pGal1(Zif268op)x6-mRuby2-tSSA1} | Kan | hygromycinR | Integrating | HO | NotI | pTEF1-ZEV driving an mRuby reporter | This study |
| pKL1-062 | {pTEF1-ZEV} | Amp | hygromycinR | Integrating | HO | NotI | Induction cassette only (no synTF) | This study |
| pKL1-067 | {Spacer} | Amp | hygromycinR | Integrating | HO | NotI | Spacer sequence for HO locus | This study |
| pKL2-181 | {pTEF1-ZEV}{pGal1(Zif268op)x6-3xFLAG-NLS-VP16-mRuby2-PDZLigS5*-tSSA1} | Kan | hygromycinR | Integrating | HO | NotI | WT synTF, no ZFTF | This study |
| pKL2-216 | {pTEF1-ZEV}{pGal1(Zif268op)x6-3xFLAG-NLS-mRuby2-42-10-WT-PDZLigS5*-tSSA1} | Kan | hygromycinR | Integrating | HO | NotI | WT synTF, no VP16 | This study |
| pKL2-255 | {pTEF1-ZEV}{pGal1(Zif268op)x6-3xFLAG-NLS-VP16-mRuby2-42-10-1x-PDZLigS5*-tSSA1} | Kan | hygromycinR | Integrating | HO | NotI | 1x R->A mutant ZF synTF | This study |
| pKL2-256 | {pTEF1-ZEV}{pGal1(Zif268op)x6-3xFLAG-NLS-VP16-mRuby2-42-10-2x-PDZLigS5*-tSSA1} | Kan | hygromycinR | Integrating | HO | NotI | 2x R->A mutant ZF synTF | This study |
| pKL2-257 | {pTEF1-ZEV}{pGal1(Zif268op)x6-3xFLAG-NLS-VP16-mRuby2-42-10-3x-PDZLigS5*-tSSA1} | Kan | hygromycinR | Integrating | HO | NotI | 3x R->A mutant ZF synTF | This study |
| pKL1-422 | {(42-10 op)x1-minCyc1-Venus-tCyc1} | Amp | -ura | Integrating | URA3 | NotI | Venus fluorescent reporter with 1x 42-10 CRMs | This study |
| pKL1-423 | {(42-10 op)x2-minCyc1-Venus-tCyc1} | Amp | -ura | Integrating | URA3 | NotI | Venus fluorescent reporter with 2x 42-10 CRMs | This study |
| pKL1-424 | {(42-10 op)x3-minCyc1-Venus-tCyc1} | Amp | -ura | Integrating | URA3 | NotI | Venus fluorescent reporter with 3x 42-10 CRMs | This study |
| pKL1-425 | {(42-10 op)x5-minCyc1-Venus-tCyc1} | Amp | -ura | Integrating | URA3 | NotI | Venus fluorescent reporter with 5x 42-10 CRMs | This study |
| pKL2-215 | {pTEF1-ZEV}{pGal1(Zif268op)x6-3xFLAG-NLS-mRuby2-42-10-4x-PDZLigS5*-tSSA1} | Kan | hygromycinR | Integrating | HO | NotI | 4x R->A mutant synTF, no VP16 | This study |
| pKL2-338 | eVOLVER mutation H345R | Kan | hygromycinR | Integrating | HO | NotI | eVOLVER mutant in clean background strain | This study |
| pKL2-339 | eVOLVER mutation A400fs(-C) | Kan | hygromycinR | Integrating | HO | NotI | eVOLVER mutant in clean background strain | This study |
| pKL2-340 | eVOLVER mutation promoter_mut | Kan | hygromycinR | Integrating | HO | NotI | eVOLVER mutant in clean background strain | This study |
| pKL2-341 | eVOLVER mutation A400fs(+C) | Kan | hygromycinR | Integrating | HO | NotI | eVOLVER mutant in clean background strain | This study |
| pKL2-342 | eVOLVER mutation promoter_mut | Kan | hygromycinR | Integrating | HO | NotI | eVOLVER mutant in clean background strain | This study |
| pKL2-343 | eVOLVER mutation A400fs(+C) | Kan | hygromycinR | Integrating | HO | NotI | eVOLVER mutant in clean background strain | This study |
| pKL2-344 | eVOLVER mutation A400fs(+C) | Kan | hygromycinR | Integrating | HO | NotI | eVOLVER mutant in clean background strain | This study |
| pKL2-345 | eVOLVER mutation A400fs(+C) | Kan | hygromycinR | Integrating | HO | NotI | eVOLVER mutant in clean background strain | This study |
| pKL2-346 | eVOLVER mutation A400fs(-C) | Kan | hygromycinR | Integrating | HO | NotI | eVOLVER mutant in clean background strain | This study |
| pKL2-347 | eVOLVER mutation A400fs(+C) | Kan | hygromycinR | Integrating | HO | NotI | eVOLVER mutant in clean background strain | This study |
| pKL2-348 | eVOLVER mutation A400fs(+C) | Kan | hygromycinR | Integrating | HO | NotI | eVOLVER mutant in clean background strain | This study |
| pKL2-349 | eVOLVER mutation G408fs(-G) | Kan | hygromycinR | Integrating | HO | NotI | eVOLVER mutant in clean background strain | This study |
| pKL2-350 | eVOLVER mutation A400fs(-C) | Kan | hygromycinR | Integrating | HO | NotI | eVOLVER mutant in clean background strain | This study |
| pKL2-351 | eVOLVER mutation A400fs(-C) | Kan | hygromycinR | Integrating | HO | NotI | eVOLVER mutant in clean background strain | This study |
| pKL2-352 | eVOLVER mutation H345Y | Kan | hygromycinR | Integrating | HO | NotI | eVOLVER mutant in clean background strain | This study |
| pKL2-353 | eVOLVER mutation T445fs(+C) | Kan | hygromycinR | Integrating | HO | NotI | eVOLVER mutant in clean background strain | This study |
| pKL2-354 | eVOLVER mutation +c in ZEV VP16 | Kan | hygromycinR | Integrating | HO | NotI | eVOLVER mutant in clean background strain | This study |
| pKL2-355 | eVOLVER mutation g -> t in Zif268 | Kan | hygromycinR | Integrating | HO | NotI | eVOLVER mutant in clean background strain | This study |
| pKL2-356 | eVOLVER mutation promoter_del | Kan | hygromycinR | Integrating | HO | NotI | eVOLVER mutant in clean background strain | This study |
| pKL2-357 | eVOLVER mutation promoter_del | Kan | hygromycinR | Integrating | HO | NotI | eVOLVER mutant in clean background strain | This study |
| pKL2-358 | eVOLVER mutation promoter_del | Kan | hygromycinR | Integrating | HO | NotI | eVOLVER mutant in clean background strain | This study |
| pKL2-359 | eVOLVER mutation A346I (frameshift = null) | Kan | hygromycinR | Integrating | HO | NotI | eVOLVER mutant in clean background strain | This study |
| pKL2-360 | eVOLVER mutation E199* | Kan | hygromycinR | Integrating | HO | NotI | eVOLVER mutant in clean background strain | This study |
| pKL2-361 | eVOLVER mutation promoter_del | Kan | hygromycinR | Integrating | HO | NotI | eVOLVER mutant in clean background strain | This study |
| pKL2-362 | eVOLVER mutation c -> g in C2H2 | Kan | hygromycinR | Integrating | HO | NotI | eVOLVER mutant in clean background strain | This study |
| pKL2-363 | eVOLVER mutation g -> c in C2H2 | Kan | hygromycinR | Integrating | HO | NotI | eVOLVER mutant in clean background strain | This study |
