## Supplementary material for "Cooperative assembly confers regulatory specificity and long-term genetic circuit stability": Table S2

**Table S2. Related to Figures 1-6.** Yeast strains used in this study

| | | Marker Loci | | |
|---|---|---|---|---|
| **Figure** | **Strain** | **HO** | **URA3** | **LEU2** |
| Fig 1B | yKL592 | pKL2-160 | pKL1-100 | pKL1-224 |
| | yKL904 | pKL1-010 | pKL1-065 | pKL1-224 |
| Fig 2B | yKL520 | pKL2-138 | pKL1-100 | pKL1-222 |
| | yKL568 | pKL2-138 | pKL1-100 | pKL1-224 |
| | yKL592 | pKL2-160 | pKL1-100 | pKL1-224 |
| | yKL948 | pKL2-255 | pKL1-100 | pKL1-224 |
| | yKL949 | pKL2-256 | pKL1-100 | pKL1-224 |
| | yKL950 | pKL2-257 | pKL1-100 | pKL1-224 |
| | yKL958 | pKL2-138 | pKL1-422 | pKL1-222 |
| | yKL959 | pKL2-138 | pKL1-423 | pKL1-419 |
| | yKL960 | pKL2-138 | pKL1-424 | pKL1-420 |
| Fig 2C | yKL520 | pKL2-138 | pKL1-100 | pKL1-222 |
| | yKL568 | pKL2-138 | pKL1-100 | pKL1-224 |
| | yKL592 | pKL2-160 | pKL1-100 | pKL1-224 |
| | yKL958 | pKL2-138 | pKL1-422 | pKL1-222 |
| | yKL950 | pKL2-257 | pKL1-100 | pKL1-224 |
| | yKL960 | pKL2-138 | pKL1-424 | pKL1-420 |
| | yKL962 | pKL2-138 | pKL1-425 | pKL1-421 |
| | yKL515 | pKL2-133 | pKL1-095 | pKL1-222 |
| | yKL518 | pKL2-136 | pKL1-098 | pKL1-222 |
| | yKL522 | pKL2-140 | pKL1-102 | pKL1-222 |
| | yKL524 | pKL2-142 | pKL1-104 | pKL1-222 |
| | yKL526 | pKL2-144 | pKL1-106 | pKL1-222 |
| | yKL529 | pKL2-147 | pKL1-109 | pKL1-222 |
| | yKL531 | pKL2-149 | pKL1-111 | pKL1-222 |
| | *yKL533* | *pKL2-151* | *pKL1-113* | *pKL1-222* |
| | yKL563 | pKL2-133 | pKL1-095 | pKL1-224 |
| | yKL566 | pKL2-136 | pKL1-098 | pKL1-224 |
| | yKL570 | pKL2-140 | pKL1-102 | pKL1-224 |
| | yKL572 | pKL2-142 | pKL1-104 | pKL1-224 |
| | yKL574 | pKL2-144 | pKL1-106 | pKL1-224 |
| | yKL577 | pKL2-147 | pKL1-109 | pKL1-224 |
| | yKL579 | pKL2-149 | pKL1-111 | pKL1-224 |
| | yKL581 | pKL2-151 | pKL1-113 | pKL1-224 |
| | yKL587 | pKL2-155 | pKL1-095 | pKL1-224 |
| | yKL590 | pKL2-158 | pKL1-098 | pKL1-224 |
| | yKL594 | pKL2-162 | pKL1-102 | pKL1-224 |
| | yKL596 | pKL2-164 | pKL1-104 | pKL1-224 |
| | yKL598 | pKL2-166 | pKL1-106 | pKL1-224 |
| | yKL601 | pKL2-169 | pKL1-109 | pKL1-224 |
| | yKL603 | pKL2-171 | pKL1-111 | pKL1-224 |
| | yKL605 | pKL2-173 | pKL1-113 | pKL1-224 |
| Fig 3A, B | yKL520 | pKL2-138 | pKL1-100 | pKL1-222 |
| | yKL568 | pKL2-138 | pKL1-100 | pKL1-224 |
| | yKL592 | pKL2-160 | pKL1-100 | pKL1-224 |
| | yKL736 | pKL1-067 | pKL1-100 | pKL1-224 |
| Fig 4B, C | yKL520 | pKL2-138 | pKL1-100 | pKL1-222 |
| | yKL568 | pKL2-138 | pKL1-100 | pKL1-224 |
| | yKL592 | pKL2-160 | pKL1-100 | pKL1-224 |
| | yKL736 | pKL1-067 | pKL1-100 | pKL1-224 |
| | yKL780 | pKL2-215 | pKL1-100 | pKL1-222 |
| | yKL788 | pKL2-215 | pKL1-100 | pKL1-224 |
| | yKL789 | pKL2-216 | pKL1-100 | pKL1-224 |
| Fig 5A | yKL520 | pKL2-138 | pKL1-100 | pKL1-222 |
| | yKL568 | pKL2-138 | pKL1-100 | pKL1-224 |
| | yKL592 | pKL2-160 | pKL1-100 | pKL1-224 |
| Fig 5B | yKL520 | pKL2-138 | pKL1-100 | pKL1-222 |
| | yKL592 | pKL2-160 | pKL1-100 | pKL1-224 |
| | yKL520 | pKL2-138 | pKL1-100 | pKL1-222 |
| | yKL568 | pKL2-138 | pKL1-100 | pKL1-224 |
| | yKL592 | pKL2-160 | pKL1-100 | pKL1-224 |
| | yKL1261 | pKL2-338 | pKL1-100 | pKL1-224 |
| | yKL1262 | pKL2-339 | pKL1-100 | pKL1-224 |
| | yKL1263 | pKL2-340 | pKL1-100 | pKL1-224 |
| | yKL1264 | pKL2-341 | pKL1-100 | pKL1-224 |
| | yKL1265 | pKL2-342 | pKL1-100 | pKL1-224 |
| | yKL1266 | pKL2-343 | pKL1-100 | pKL1-224 |
| | yKL1267 | pKL2-344 | pKL1-100 | pKL1-224 |
| | yKL1268 | pKL2-345 | pKL1-100 | pKL1-224 |
| | yKL1269 | pKL2-346 | pKL1-100 | pKL1-224 |
| | yKL1270 | pKL2-347 | pKL1-100 | pKL1-224 |
| | yKL1271 | pKL2-348 | pKL1-100 | pKL1-224 |

| | | | | |
|---|---|---|---|---|
| Fig 5C | yKL1272 | pKL2-349 | pKL1-100 | pKL1-224 |
| | yKL1273 | pKL2-350 | pKL1-100 | pKL1-224 |
| | yKL1274 | pKL2-351 | pKL1-100 | pKL1-224 |
| | yKL1275 | pKL2-352 | pKL1-100 | pKL1-224 |
| | yKL1276 | pKL2-353 | pKL1-100 | pKL1-224 |
| | yKL1277 | pKL2-354 | pKL1-100 | pKL1-224 |
| | yKL1278 | pKL2-355 | pKL1-100 | pKL1-224 |
| | yKL1279 | pKL2-356 | pKL1-100 | pKL1-224 |
| | yKL1280 | pKL2-357 | pKL1-100 | pKL1-224 |
| | yKL1281 | pKL2-358 | pKL1-100 | pKL1-224 |
| | yKL1282 | pKL2-359 | pKL1-100 | pKL1-224 |
| | yKL1283 | pKL2-360 | pKL1-100 | pKL1-224 |
| | yKL1284 | pKL2-361 | pKL1-100 | pKL1-224 |
| | yKL1285 | pKL2-362 | pKL1-100 | pKL1-224 |
| | yKL1286 | pKL2-363 | pKL1-100 | pKL1-224 |
| Fig 5D | yKL592 | pKL2-160 | pKL1-100 | pKL1-224 |
| | yKL736 | pKL1-067 | pKL1-100 | pKL1-224 |
| | yKL1268 | pKL2-345 | pKL1-100 | pKL1-224 |
| | yKL1285 | pKL2-362 | pKL1-100 | pKL1-224 |
| Fig 5E | yKL520 | pKL2-138 | pKL1-100 | pKL1-222 |
| | yKL568 | pKL2-138 | pKL1-100 | pKL1-224 |
| | yKL592 | pKL2-160 | pKL1-100 | pKL1-224 |
| | yKL736 | pKL1-067 | pKL1-100 | pKL1-224 |
| | yKL1268 | pKL2-345 | pKL1-100 | pKL1-224 |
| | yKL1285 | pKL2-362 | pKL1-100 | pKL1-224 |
| Fig S1B | yKL044 | pKL2-020 | | |
| Fig S1D | yKL520 | pKL2-138 | pKL1-100 | pKL1-222 |
| | yKL568 | pKL2-138 | pKL1-100 | pKL1-224 |
| | yKL592 | pKL2-160 | pKL1-100 | pKL1-224 |
| | yKL732 | pKL1-062 | pKL1-100 | pKL1-222 |
| | yKL736 | pKL1-067 | pKL1-100 | pKL1-224 |
| | yKL737 | pKL1-062 | pKL1-100 | pKL1-224 |
| | yKL738 | pKL2-181 | pKL1-100 | pKL1-224 |
| | yKL789 | pKL2-216 | pKL1-100 | pKL1-224 |
| Fig S1E | yKL515 | pKL2-133 | pKL1-095 | pKL1-222 |
| | yKL516 | pKL2-134 | pKL1-096 | pKL1-222 |
| | yKL517 | pKL2-135 | pKL1-097 | pKL1-222 |
| | yKL518 | pKL2-136 | pKL1-098 | pKL1-222 |
| | yKL519 | pKL2-137 | pKL1-099 | pKL1-222 |
| | yKL520 | pKL2-138 | pKL1-100 | pKL1-222 |
| | yKL521 | pKL2-139 | pKL1-101 | pKL1-222 |
| | yKL522 | pKL2-140 | pKL1-102 | pKL1-222 |
| | yKL523 | pKL2-141 | pKL1-103 | pKL1-222 |
| | yKL524 | pKL2-142 | pKL1-104 | pKL1-222 |
| | yKL525 | pKL2-143 | pKL1-105 | pKL1-222 |
| | yKL526 | pKL2-144 | pKL1-106 | pKL1-222 |
| | yKL527 | pKL2-145 | pKL1-107 | pKL1-222 |
| | yKL528 | pKL2-146 | pKL1-108 | pKL1-222 |
| | yKL529 | pKL2-147 | pKL1-109 | pKL1-222 |
| | yKL530 | pKL2-148 | pKL1-110 | pKL1-222 |
| | yKL531 | pKL2-149 | pKL1-111 | pKL1-222 |
| | yKL532 | pKL2-150 | pKL1-112 | pKL1-222 |
| | *yKL533* | *pKL2-151* | *pKL1-113* | *pKL1-222* |
| | yKL536 | pKL2-154 | pKL1-116 | pKL1-222 |
| | yKL563 | pKL2-133 | pKL1-095 | pKL1-224 |
| | yKL564 | pKL2-134 | pKL1-096 | pKL1-224 |
| | yKL565 | pKL2-135 | pKL1-097 | pKL1-224 |
| | yKL566 | pKL2-136 | pKL1-098 | pKL1-224 |
| | yKL567 | pKL2-137 | pKL1-099 | pKL1-224 |
| | yKL568 | pKL2-138 | pKL1-100 | pKL1-224 |
| | yKL569 | pKL2-139 | pKL1-101 | pKL1-224 |
| | yKL570 | pKL2-140 | pKL1-102 | pKL1-224 |
| | yKL571 | pKL2-141 | pKL1-103 | pKL1-224 |
| | yKL572 | pKL2-142 | pKL1-104 | pKL1-224 |
| | yKL573 | pKL2-143 | pKL1-105 | pKL1-224 |
| | yKL574 | pKL2-144 | pKL1-106 | pKL1-224 |
| | yKL575 | pKL2-145 | pKL1-107 | pKL1-224 |
| | yKL576 | pKL2-146 | pKL1-108 | pKL1-224 |
| | yKL577 | pKL2-147 | pKL1-109 | pKL1-224 |
| | yKL578 | pKL2-148 | pKL1-110 | pKL1-224 |
| | yKL579 | pKL2-149 | pKL1-111 | pKL1-224 |
| | yKL580 | pKL2-150 | pKL1-112 | pKL1-224 |
| | yKL581 | pKL2-151 | pKL1-113 | pKL1-224 |
| | yKL584 | pKL2-154 | pKL1-116 | pKL1-224 |
| | yKL587 | pKL2-155 | pKL1-095 | pKL1-224 |
| | yKL588 | pKL2-156 | pKL1-096 | pKL1-224 |
| | yKL589 | pKL2-157 | pKL1-097 | pKL1-224 |

| | | | | |
|---|---|---|---|---|
| | yKL590 | pKL2-158 | pKL1-098 | pKL1-224 |
| | yKL591 | pKL2-159 | pKL1-099 | pKL1-224 |
| | yKL593 | pKL2-161 | pKL1-101 | pKL1-224 |
| | yKL594 | pKL2-162 | pKL1-102 | pKL1-224 |
| | yKL595 | pKL2-163 | pKL1-103 | pKL1-224 |
| | yKL596 | pKL2-164 | pKL1-104 | pKL1-224 |
| | yKL597 | pKL2-165 | pKL1-105 | pKL1-224 |
| | yKL598 | pKL2-166 | pKL1-106 | pKL1-224 |
| | yKL599 | pKL2-167 | pKL1-107 | pKL1-224 |
| | yKL600 | pKL2-168 | pKL1-108 | pKL1-224 |
| | yKL601 | pKL2-169 | pKL1-109 | pKL1-224 |
| | yKL602 | pKL2-170 | pKL1-110 | pKL1-224 |
| | yKL603 | pKL2-171 | pKL1-111 | pKL1-224 |
| | yKL604 | pKL2-172 | pKL1-112 | pKL1-224 |
| | yKL605 | pKL2-173 | pKL1-113 | pKL1-224 |
| | yKL608 | pKL2-176 | pKL1-116 | pKL1-224 |
| Fig S3B | yKL520 | pKL2-138 | pKL1-100 | pKL1-222 |
| | yKL950 | pKL2-257 | pKL1-100 | pKL1-224 |
| Fig S3C | yKL520 | pKL2-138 | pKL1-100 | pKL1-222 |
| | yKL568 | pKL2-138 | pKL1-100 | pKL1-224 |
| | yKL592 | pKL2-160 | pKL1-100 | pKL1-224 |
| | yKL736 | pKL1-067 | pKL1-100 | pKL1-224 |
| Fig S4A-E | yKL520 | pKL2-138 | pKL1-100 | pKL1-222 |
| | yKL568 | pKL2-138 | pKL1-100 | pKL1-224 |
| | yKL592 | pKL2-160 | pKL1-100 | pKL1-224 |
| | yKL736 | pKL1-067 | pKL1-100 | pKL1-224 |
| Fig S5A, C | yKL520 | pKL2-138 | pKL1-100 | pKL1-222 |
| | yKL568 | pKL2-138 | pKL1-100 | pKL1-224 |
| | yKL592 | pKL2-160 | pKL1-100 | pKL1-224 |
| | yKL736 | pKL1-067 | pKL1-100 | pKL1-224 |
| Fig S5B,D,E | yKL520 | pKL2-138 | pKL1-100 | pKL1-222 |
| | yKL568 | pKL2-138 | pKL1-100 | pKL1-224 |
| | yKL592 | pKL2-160 | pKL1-100 | pKL1-224 |
| | yKL736 | pKL1-067 | pKL1-100 | pKL1-224 |
| | yKL780 | pKL2-215 | pKL1-100 | pKL1-222 |
| | yKL788 | pKL2-215 | pKL1-100 | pKL1-224 |
| | yKL789 | pKL2-216 | pKL1-100 | pKL1-224 |
| Fig S6A | yKL520 | pKL2-138 | pKL1-100 | pKL1-222 |
| | yKL568 | pKL2-138 | pKL1-100 | pKL1-224 |
| | yKL592 | pKL2-160 | pKL1-100 | pKL1-224 |
| | yKL619 | pKL1-067 | pKL1-065 | pKL1-224 |
| | yKL736 | pKL1-067 | pKL1-100 | pKL1-224 |
| Fig S6B | yKL1261 | pKL2-338 | pKL1-100 | pKL1-224 |
| | yKL1262 | pKL2-339 | pKL1-100 | pKL1-224 |
| | yKL1263 | pKL2-340 | pKL1-100 | pKL1-224 |
| | yKL1264 | pKL2-341 | pKL1-100 | pKL1-224 |
| | yKL1265 | pKL2-342 | pKL1-100 | pKL1-224 |
| | yKL1266 | pKL2-343 | pKL1-100 | pKL1-224 |
| | yKL1267 | pKL2-344 | pKL1-100 | pKL1-224 |
| | yKL1268 | pKL2-345 | pKL1-100 | pKL1-224 |
| | yKL1269 | pKL2-346 | pKL1-100 | pKL1-224 |
| | yKL1270 | pKL2-347 | pKL1-100 | pKL1-224 |
| | yKL1271 | pKL2-348 | pKL1-100 | pKL1-224 |
| | yKL1272 | pKL2-349 | pKL1-100 | pKL1-224 |
| | yKL1273 | pKL2-350 | pKL1-100 | pKL1-224 |
| | yKL1274 | pKL2-351 | pKL1-100 | pKL1-224 |
| | yKL1275 | pKL2-352 | pKL1-100 | pKL1-224 |
| | yKL1276 | pKL2-353 | pKL1-100 | pKL1-224 |
| | yKL1277 | pKL2-354 | pKL1-100 | pKL1-224 |
| | yKL1278 | pKL2-355 | pKL1-100 | pKL1-224 |
| | yKL1279 | pKL2-356 | pKL1-100 | pKL1-224 |
| | yKL1280 | pKL2-357 | pKL1-100 | pKL1-224 |
| | yKL1281 | pKL2-358 | pKL1-100 | pKL1-224 |
| | yKL1282 | pKL2-359 | pKL1-100 | pKL1-224 |
| | yKL1283 | pKL2-360 | pKL1-100 | pKL1-224 |
| | yKL1284 | pKL2-361 | pKL1-100 | pKL1-224 |
| | yKL1285 | pKL2-362 | pKL1-100 | pKL1-224 |
| | yKL1286 | pKL2-363 | pKL1-100 | pKL1-224 |
| Fig S6C | yKL515 | pKL2-133 | pKL1-095 | pKL1-222 |
| | yKL563 | pKL2-133 | pKL1-095 | pKL1-224 |
| | yKL587 | pKL2-155 | pKL1-095 | pKL1-224 |
| | yKL623 | pKL1-067 | pKL1-095 | pKL1-224 |
| | yKL907 | pKL2-020 | pKL1-065 | pKL1-224 |
| Fig S6D | yKL515 | pKL2-133 | pKL1-095 | pKL1-222 |
| | yKL587 | pKL2-155 | pKL1-095 | pKL1-224 |
